# Dietary Microplastics Engage Gut Mechanosensory-Endocrine Signaling and Disrupt Bone Homeostasis

**DOI:** 10.64898/2026.04.03.716216

**Authors:** Aaron S. Romero, Sumira Phatak, Sanchiti Patil, Hamid Y. Dar, Jaclyn A. Rivas, Olufunmilola M Oyebamiji, Brianna B. Maes, Siem S. Goitom, Crystal Madera Enriquez, Jazmin Orozco, Cristina N. Coffman, Ruixuan Liu, Julie G. In, Matthew J. Campen, Kel Cook, Richard Levenson, Jessica M. Gross, Shuguang Leng, Anthony Cretara, Roberto Pacifici, Eliseo F. Castillo

## Abstract

**Background and Aims:** Microplastics are pervasive environmental contaminants increasingly detected in food and water supplies; however, their effects on gastrointestinal physiology and systemic health remain poorly understood. We investigated whether chronic dietary microplastic exposure alters colonic neuroendocrine signaling and skeletal health.

**Methods:** Female and male C57BL/6J mice were fed purified basal, high-fat/high-cholesterol, or high-fiber diets with or without a physiological relevant polystyrene microplastic mixture (∼1.7 mg/kg; particle sizes 0.49 – 5.0 µm) for 12 weeks. Colonic cellular responses were evaluated using ELISA, histology, immunofluorescence, and single-nuclei RNA sequencing. Fecal microbiota transplantation was performed to assess microbial contributions to microplastic-induced phenotypes. Bone microarchitecture was assessed by micro-computed tomography. Human bone specimens were analyzed for microplastic content, and primary osteoblast mineralization assays were performed.

**Results:** Dietary microplastic exposure increased chromogranin A-positive enteroendocrine cells and enhanced serotonergic signaling in the colon without evidence of intestinal inflammation or lineage reprogramming. Single-nuclei transcriptomic analysis identified compartment-specific serotonergic and mechanosensory adaptations in epithelial and enteric neuronal populations. Transfer of microbiota from microplastic-exposed donors to control recipients recapitulated increased enteroendocrine cell abundance. Chronic microplastic ingestion induced sex- and diet-dependent reductions in trabecular bone loss and architecture without systemic inflammatory activation. Microplastics were detected in human mineralized bone, and microplastic exposure impaired osteoblast mineralization in a donor-dependent manner.

**Conclusions:** Chronic ingestion of microplastics remodels gut neuroendocrine signaling through microbiota-dependent mechanisms and impairs skeletal homeostasis in the absence of overt inflammation. These findings identify a previously unrecognized gut-bone pathway through which dietary microplastic exposure may influence host physiology.

## Introduction

Over the past few decades, MPs have emerged as pervasive environmental contaminants^1, 2^, raising growing concern about their impact on human health^3^. Most existing studies linking MP to disease are correlative, focusing on MP detection in diseased tissues^4–7^ or have noted altered tissue and cellular metabolism^8–12^. Ingestion of MP is the major route of exposure making the gastrointestinal (GI) tract the first point of contact. Interestingly, there are numerous studies linking MP exposure to intestinal inflammation as well as exacerbating intestinal inflammation in various colitis models ^13–17^. MPs have also been shown to alter the gut microbiota and microbial metabolites ^8, 18^. In addition, MPs are considered endocrine disruptors with studies showing effects on various endocrine organs such as the adrenal gland, ovaries and thyroid ^19–24^. These interactions and effects are particularly relevant in light of growing evidence that enteroendocrine cells (EECs) represent an early interface between luminal content and host metabolism and physiology. As the gut’s hormone-producing sentinels, these cells integrate luminal cues with systemic hormone signaling. A recent study showed MP exposure during a chronic colitis model induced by dextran sulfate sodium resulted in increased levels of chromogranin A-positive cells, a marker of EECs, in the colon^25^.

Given the central role of EEC signals in systemic physiology, these observations raise the possibility the MP-induced alterations in the gut may extend to distal tissues such as the skeletal system. EEC-derived hormones such as serotonin can influence bone homeostasis^26–29^. There have been studies showing MPs alter the skeletal compartment. Daily oral gastric gavage (OGG) of 5-µm polystyrene (PS)-MPs or modified PS-MPs for 28 days in mice impaired trabecular microarchitecture. Interestingly, PS-MPs accumulated in bones, potentially within the marrow compartment, although the exact localization was not determined^30^. Accumulation of 80 nm-sized PS nanoplastics (PSNP) were also found in bone tissue after daily OGG for 42 days, which resulted in impaired hematopoiesis in both the bone marrow and spleen, and reduced hematopoietic stem/progenitor cell populations. Mechanistically, PSNP exposure induced oxidative stress and activated oxeiptosis and senescence pathways, contributing to hematotoxicity^31^. Similar effects on the bone marrow were also observed after daily OGG of 5- and 10-µm sized for 42 days^32^. In rats, daily OGG of 1-µm PS MP impaired endochondral ossification by disrupting chondrocyte organization and inducing ER stress^33^. Another study showed intraperitoneal injection into rats caused skeletal toxicity^34^. These studies support a growing link between MP exposure and skeletal dysfunction primarily through systemic inflammation or marrow toxicity. Nevertheless, the precise impact of MPs on the gut-bone axis remains to be fully elucidated.

A major caveat with these prior *in vivo* studies linking MP to intestinal inflammation or bone loss is that they predominantly relied on bolus oral gavage, intraperitoneal injection, or delivery in drinking water ^25, 30–34^. These exposure paradigms may not fully recapitulate typical dietary intake including regulated ingestion, normal digestive processing, and sustained luminal interactions within the gut. Dietary exposure may closely reflect human consumption by integrating MPs into physiological contexts that involve GI motility, mucosal contact, and GI hormone release, all of which could influence MP uptake and downstream systemic effects. Diet might plausibly exacerbate or mitigate MP toxicity: high-fat, high-cholesterol diet could enhance MP absorption through impaired barrier integrity or enhanced lipid-mediated transport, whereas a high-fiber diet may limit MP uptake by strengthening mucosal defenses and reinforcing epithelial barrier function. Although diet is a well-established regulator of the gut function, metabolism, and endocrine signaling, its role in shaping MP-induced toxicity, particularly with respect to systemic physiology and underlying causal mechanisms, remains poorly understood. Herein, we demonstrate that chronic dietary MP exposure alters colonic EEC and serotonin signaling, partially mediated by the gut microbiota, while also disrupting skeletal homeostasis.

## Materials and methods

### Animals

Male and female C57BL/6J (Strain #000664) were purchased from The Jackson Laboratory (JAX) at three weeks of age. Additionally, male and female B6(SJL)-*Piezo2*^tm1.1(cre)Apat^/J (Strain #027719) PIEZO2-GFP reporter mice were purchased from JAX between 5-7 weeks of age. Animals were maintained at constant temperature (20-24°C), relative humidity (30%-60%), and 12-hlight/dark cycle throughout the study. Animals were provided with multiple diets and water ad libitum. All experiments were approved by the Institutional Animal Care and Use Committee of the University of New Mexico Health Sciences Center, in accordance with the National Institutes of Health guidelines for use of live animals. The University of New Mexico Health Sciences Center is accredited by the American Association of Accreditation of Laboratory Animal Care.

### Dietary Exposure to Microplastics

Mice were exposed to a mixture of polystyrene microspheres via their diet over a 12-wk period with either 0% or 1.697% polystyrene microspheres (Magsphere, Pasadena, CA). Exposure was based on an estimated average human intake of 5 grams plastics ingested per week^35^. Five sizes of polystyrene microspheres were utilized by mass: 0.49 µm (Magsphere, catalog no. PSFB500NM), 1.0 µm (Magsphere, catalog no. PSMG001UM), 1.9 µm (Magsphere, catalog no. PSOF002UM), 3.1 µm (Magsphere, catalog no. PSFY003UM), and 5.0 µm (Magsphere, catalog no. PSFR005UM) (**Supplementary Table 1**). Base diets included AIN-93M (AIN, Product no. D10012MO), 40% fat from lard plus 1.25% added cholesterol (HFC, Product no. D24020804), and 10% added inulin high-fiber (FIB, Product no. D24020806) purified formulas (Research Diets Inc., New Brunswick, NJ) (**Supplementary Table 2**). Microspheres were washed 10 times using sterile, deionized water to remove surfactant and sodium azide prior to formulation with AIN (AMP, Product no. D24020803), HFC (HMP, Product no. D24020805), and FIB (FMP, Product no. D24020807) diets. Mice were weighed weekly and food intake was tracked biweekly throughout the study. After 12 weeks, mice were euthanized using isoflurane and exsanguinated, then systemically perfused with ice cold saline to ensure removal of blood from major organs.

### Cells and Tissue Preparation

Upon necropsy, tissues were weighed, measured, sectioned out for various procedures, and flash frozen. Samples collected for downstream analysis included colon, liver, plasma, spinal cords, femurs, and stool. These samples were either flash frozen in liquid nitrogen and stored in −80C, fixed with formalin, or put into 70% ethanol (EtOH). Blood was collected via cardiac puncture and collected in the BD P800 blood collection tube containing proprietary cocktail of protease, esterase and DPP-IV Inhibitors (BD Biosciences, catalog no. 366420) for plasma isolation. Plasma was used for serotonin (5-HT) detection via ELISA (Abnova, Catalog no. KA1894) or hormone (Amylin; C-Peptide; Ghrelin; GIP; GLP-1 (total); Glucagon; Insulin; Leptin; PP; PYY; Resistin; and Secretin) and cytokine (IL-6, MCP-1/CCL2; TNFα) detection by a MILLIPLEX mouse metabolic hormone expanded panel multiplex assay (Millipore, catalog no. MMHE-44K). Stool samples collected from weeks 0, 3, 6, 9, and 12 were processed and normalized to 90mg/mL using PBS + 0.1% TWEEN to analyze inflammatory markers of the colon such as Lipocalin-2 (LCN2)^36^ using R&D Systems Quantikine ELISA Mouse Lipocalin-2/NGAL Immunoassay (Catalog no. MLCN20) and secretory IgA (sIgA) from Bio-Techne Mouse Secretory IgA ELISA Kit (colorimetric) (Catalog no. NBP3-11824). A section of the proximal colon was stored in 10% formalin for 48 hours, then washed and stored in 70% EtOH prior to processing for tissue sectioning. Histopathological evaluation was performed on mouse colon tissue. Formalin-fixed, paraffin-embedded (FFPE) sections of proximal colon were prepared at the University of New Mexico Comprehensive Cancer Center (UNMCCC) Human Tissue Repository (HTR) Core Facility. Sections (5 μm) were cut from blocks fixed in 10% neutral-buffered formalin and stained with hematoxylin and eosin (H&E) using standard protocols. The degree of intestinal inflammation scoring was performed based on the guidelines provided by Erben et al^37^. Colon sections were scored for histopathological features of colitis based on established criteria, including: submucosal edema (0–3), polymorphonuclear leukocyte (PMN) infiltration (0–3), goblet cell depletion (0–3), and epithelial integrity (0–3) (**Supplementary Table 3**). Liver sections were evaluated using the NASH Clinical Research Network Scoring System following the guidelines provided by Liang et al ^38^. Assessed histopathological parameters included - macrovesicular steatosis (+/−), microvesicular steatosis (+/−), steatosis grade (0–3), lobular inflammation (0–3), hepatocellular hypertrophy (0–2), and fibrosis stage (0–4) (**Supplementary Table 4**).

### Fecal Microbiota Transplant (FMT)

Fresh stool pellets were collected from several C57BL/6J mice that were either exposed to microplastics or no microplastics in their diet. Pellets were weighed, standardized, and then resuspended in sterile 1xPBS. The PBS fecal slurry was filtered and then used for FMT in two groups of PIEZO2-GFP reporter mice pretreated with antibiotics as we have previously described^36^. These mice received water supplemented with Baytril (Enrofloxacinoral, 100 mg/mL). After one week of antibiotic treatment, two groups of male PIEZO2-GFP reporter mice were given a slurry of bacteria (90mg/mL) via oral gastric gavage from C57BL/6J mice that were either treated with or without microplastics in their diet. Twelve-weeks after FMT, mouse colons and bones were collected for further assessment.

### Micro-CT Analysis

Spinal cords and femurs were collected from all mice to measure cortical area and thickness as well as trabecular volume and separation using a micro-CT (µCT) scanner as we have previously reported ^39, 40^. Indices of trabecular and cortical bone volume and structure were measured in the spine (excised 5th lumbar spine body) and the femur, respectively, using a Scanco mCT-40 scanner (Scanco Medical, Bassersdorf, Switzerland). µCT scanning and analysis was performed as reported previously. Briefly, trabecular and cortical bone regions were evaluated using isotropic 12-mm voxels. For the vertebral trabecular region, we evaluated 250 transverse CT slices between the cranial and caudal end plates, excluding 100mm near each end plate. For the femoral trabecular region, we analyzed 100 slices from the 50 slices under the distal growth plate. Femoral cortical bone was assessed using 50 continuous CT slides located at the femoral midshaft.

### Generation of intestinal organoids

Intestinal organoid media was comprised of Advanced Dulbecco’s modified Eagle medium/Ham’s F-12 (ThermoFisher, Waltham, MA), 100 U/mL penicillin/streptomycin (Quality Biological, Gaithersburg, MD), WNT surrogate-Fc fusion protein (ImmunoPrecise Antibodies Ltd, Utrecht, The Netherlands) 15% v/v R-spondin1 conditioned medium (cell line kindly provided by Calvin Kuo, Stanford University), 10% v/v Noggin conditioned medium (cell line kindly provided by Gijs van den Brink, Tytgat Institute for Liver and Intestinal Research), 1X B27 supplement (ThermoFisher), 10 mM HEPES (ThermoFisher), 1X GlutaMAX (ThermoFisher), 1mM N-acetylcysteine (MilliporeSigma), 50 ng/mL human epidermal growth factor (ThermoFisher), 10 nM [Leu-15] gastrin (AnaSpec, Fremont, CA), 500 nM A83-01 (Tocris, Bristol, United Kingdom), 10 μM SB202190 (MilliporeSigma), 100 mg/mL primocin (InvivoGen, San Diego, CA). Base media, for differentiation of organoids, had the same composition of media but lacked WNT surrogate Fc fusion protein, Rspo-1 and SB202190. Colonic crypt isolation and colonic organoid generation were prepared as previously reported ^41–44^. Isolated crypts from proximal mouse colon were resuspended in Matrigel (Corning) and 25 µL droplets were plated in a 24-well tissue culture plate (Corning). After polymerization at 37°C, 500µL of organoid expansion media was added for 2 d. After 2 d, the organoid expansion media was replaced every other day. Colonic organoids were passaged every 2-3 days by harvesting in Cultrex Organoid Harvesting Solution (Bio-Techne, Minneapolis, MN) at 4°C with shaking for 45 min, as previously described ^43, 44^. All colonic organoids cultures were maintained at 37°C and 5% CO2. Unless noted, colonic organoids lines have been passaged >30 times. Colonic organoids were harvested from Matrigel using Cultrex Organoid Harvesting Solution as previously described ^43, 44^.

### RNA Isolation, Quantification, and RT-qPCR

RNA Isolation was performed on C57BL/6J mice femurs following shaving of muscle off bone, mashing of the bone with a mortar and pestle, and being put into TRIzol Reagent (Thermofisher) then using the RNA Purelink Minikit (Thermofisher) according to the manufacturer’s protocol. RNA isolation was performed using the RNA Purelink Minikit (Invitrogen) according to the manufacturer’s protocols on colonic organoids which were harvested from Matrigel using Cultrex Organoid Harvesting Solution as previously described^44^. RNA was quantified using a Nanodrop2000 and all samples yielded a 260/280 of 2 ± 0.15. cDNA synthesis was performed using a BioRad MyCycler Thermal Cycler and SuperScript™ IV VILO™ Master Mix (Thermofisher) and normalizing to the lowest yielding RNA concentrations. TaqMan™ Universal PCR Master Mix (Thermofisher) was used to perform qPCR using primers listed in **Supplementary Table 5**. Using 384 well plates, qPCR was performed on a CFX384 Touch Real-Time PCR Detection System.

### Microscopy, Image analysis, Immunofluorescence and Histology

Unstained slides of paraffin-embedded colon and livers from mice or human femurs (**Supplementary Table 6**, OriGene Technologies, Inc) were stained with antibodies or dyes listed (**Supplementary Table 7**) using antigen retrieval. Briefly, slides were baked for 2 hours and allowed to cool for 30 mins. Slides were subsequently washed with xylene 2x and EtOH 5x at 100, 95, 80, 70, and 50% concentrations. Slides were then placed in 10mM of sodium citrate buffer (2.94g/1L of H2O) and microwaved on high for 5 minutes following by cooling at RT for 10 minutes 2x. After the second buffer placement, slides were placed in fresh sodium citrate buffer for 20 minutes to cool. Slides were then immersed in 0.85% NaCl for 5 minutes, dipped in ddH2O 3x and washed with 0.01M Dulbecco’s PBS 2x. Slides then had PAP pen circling and were allowed to dry for 15 minutes and washed with PBS 2x. Slides were then incubated with Image-iT FX Signal Enhancer in a H2O humidity chamber for 30 mins, and placed in primary antibody overnight in perm-block buffer (0.1% saponin, 2% BSA, 15% FBS, brought up in 1X PBS). After overnight incubation, slides were washed with 0.01M PBS 3x and incubated with secondary antibody for 2 hours in immuno-block buffer. Slides were washed 3x with PBS, 1x with ddH2O, and dried for 30 minutes. Slides then had a coverslip mounted with fructose-glycerol solution (2.5M fructose-60% glycerol stock). Immunofluorescence images were captured utilizing a Zeiss LSM800 Confocal Microscope at 40X objectives. Images were processed using Zeiss ZEN (Version 3.7) software. Imaging of H&E-stained and MP-Dye tissue sections was performed using the Evident APEXVIEW APX100 Benchtop Fluorescence Microscope, which supports both brightfield and fluorescence imaging with appropriate filter sets (e.g., DAPI, FITC, TRITC). Acquisition parameters in the cellSens APEX software were standardized across all samples, and images were exported as high-resolution TIFF files. All primary and secondary antibodies used for confocal staining are listed in **Supplementary Table 7**.

### Single-nuclei RNA Sequencing and Bioinformatic Analysis

Colonic tissue from female mice maintained on AIN or AMP diets (n = 4 biological replicates per group) was enzymatically dissociated using the Chromium Nuclei Isolation kit with RNase Inhibitor (Catalog no. PN-1000494, 10x Genomics) to generate single-nuclei suspensions. Single-nuclei RNAseq libraries were prepared using Chromium Next GEM 3’ Reagent Kit v3.1 following manufacturer’s instruction (protocol CG000315 Rev F, 10x Genomics). In summary, single-nuclei suspensions were combined with a master mix, loaded into microfluidic chips, and partitioned into gel beads-in-emulsion (GEMs). RNAs were captured and cDNAs made with primers containing a poly(dT) sequence, unique molecular identifier (UMI), GEM-specific barcode, and Illumina TruSeq adapter. After cDNA synthesis, GEMs were broken, cDNAs were amplified with 13 PCR cycles, and libraries were prepared from the amplified cDNA. After library construction, an additional 7-cycle PCR step was performed to replace Illumina sequencing adapters, and libraries were sequenced on the Singular Genomics G4 platform targeting at least 10,000 paired-end reads per nucleus. Data Processing and Quality Control - raw base call files were processed using Cell Ranger (v9.0.1) against the mm10 reference genome to generate gene-cell count matrices. Data Processing and Quality Control. Downstream analyses were performed in R (v4.4.3) using Seurat (v5.3.0) ^45^. Ambient RNA contamination was corrected using SoupX^46^ prior to filtering. Doublets were identified and removed using scDblFinder^47^. Cells were excluded based on the following criteria: <395 detected genes; >4716 detected genes; >15% mitochondrial transcripts. After filtering, 35,041 high-quality cells were retained for downstream analysis. Normalization and variance stabilization were performed using SCTransform with regression of mitochondrial percentage and sequencing depth. Samples were integrated using Harmony^48^ to correct for batch effects across biological replicates. Principal component analysis (PCA) was performed on the integrated object, and significant components were selected based on elbow plot inspection and variance explained. UMAP was used for visualization. Graph-based clustering was performed using Seurat’s FindNeighbors and FindClusters functions (resolution = 0.6). Cell types were annotated based on established canonical marker genes and cluster-specific differentially expressed genes. To preserve biological replication and avoid pseudoreplication, differential expression testing was performed using a pseudobulk approach implemented in muscat^49^. For each annotated cell type, raw counts were aggregated per biological replicate. Statistical testing was performed treating each animal as the unit of replication using edgeR/DESeq2-based modeling, with diet as the primary covariate. Multiple testing correction was performed using the Benjamini-Hochberg method. Genes with FDR-adjusted P < 0.05 were considered statistically significant. Differential abundance of cell populations was assessed by comparing the relative frequency of each annotated cell type per biological replicate between diets using Speckle ^50^. P values < 0.05 were considered significant. Raw and processed sequencing data have been deposited in GEO under accession number (will be provided upon acceptances). No regression of biological covariates (e.g., diet) was performed during integration to avoid removal of true biological signal.

### Osteoblast Experiment

Three independent human osteoblast cell lines (PromoCell, Catalog no. C-12720, **Supplementary Table 8**) were selected, all harvested from the femoral head of Caucasian non-smokers with osteoarthrosis but not osteoporosis: male age 74 (lot no. 519Z062), female age 72 (lot no. 501Z014.2), and female age 64 (lot no. 498Z010.1). Upon confluency in Osteoblast Growth Medium, each cell line was treated with either 10 ng/mL of 5-HT (BioTechne, Catalog no. 3547/50), 50 ng/mL of 5-HT, 1 µg/mL of 1 µm polystyrene (PS) microspheres (Magsphere, Catalog no. PS001UM), 1 µg/mL of 5 µm PS microspheres (Magsphere, Catalog no. PS005UM), or vehicle control for 10 days in Osteoblast Growth Medium. Following this initial exposure, cells were maintained under the same treatment conditions in Osteoblast Mineralization Medium for an additional 21 days. At the conclusion of the treatment period, mineral deposition was assessed by staining cells with Alizarin Red S (pH 4.1) for 30 minutes at room temperature, protected from light. Cells were subsequently washed three times with Dulbecco’s phosphate-buffered saline (PBS) without Ca²⁺ or Mg²⁺, then counterstained with DAPI. Fluorescence intensity was quantified using a Cytation™ 5 plate reader at a resolution of 15×15 reads per well (alizarin *λ_ex_* = 496 nm and *λ_e_*_m_ = 616 nm; DAPI *λ_ex_* = 359 nm and *λ_e_*_m_ = 451 nm) to determine osteoblast mineralization relative to nuclei count.

### Statistical Analysis

Statistical analysis was performed as described in figure legends and graphs generated display mean (±SD or SEM) and were obtained using GraphPad Prism software. If female and male data were analyzed separately, open circles, squares and triangles represent female mice whereas closed circles, squares and triangles represent male mice. The data were analyzed using two-tailed unpaired Student’s t test, a two-way ANOVA with Tukey’s multiple comparison test or described in the figure legend (GraphPad Prism v10.6.1 (892)).

## Results

### Dietary MP Exposure Alters Body Composition and Bone Mineral Density in a Sex-Dependent Manner

We set out to examine the influence of MPs in various purified formulas and the consequences they have on colonic and systemic tissues. Adolescent C57BL/6J (B6) female and male mice were fed one of three diets (detailed in **Supplementary Table 2**): (i) basal diets (modified AIN-93M, used as the base for all groups; herein called AIN), (ii) a high-fat, high-cholesterol diet (40% lard-base fat and 1.25% cholesterol; herein called HFC), or (iii) a high-fiber diet (10% added cellulose and inulin; herein called FIB) at 3 weeks of age to acclimate to dietary formulas. Female and male mice began dietary interventions at 4 weeks of age and were monitored for 12 weeks (**Figure 1A**). Each diet was provided with or without a physiologically relevant mixture of PS MPs (∼0.49, 1.0, 1.9, 3.1, and 5.0 µm; ∼1.7 mg/kg) (**Supplementary Table 1**). Diets with MPs are referred to as AMP (AIN with MPs), HMP (HFC with MPs), and FMP (FIB with MPs). Across all dietary groups, endpoint weight gain was unaltered between AIN and AMP groups, HFC and HMP groups, and FIB and FMP groups (**Supplementary Fig. 1A-F**). Similarly, food (**Supplementary Fig. 1G-L**) and energy (**Supplementary Fig. 1M-R**) intake remained consistent, allowing direct comparison of weight and body composition outcomes.

**Figure 1.**
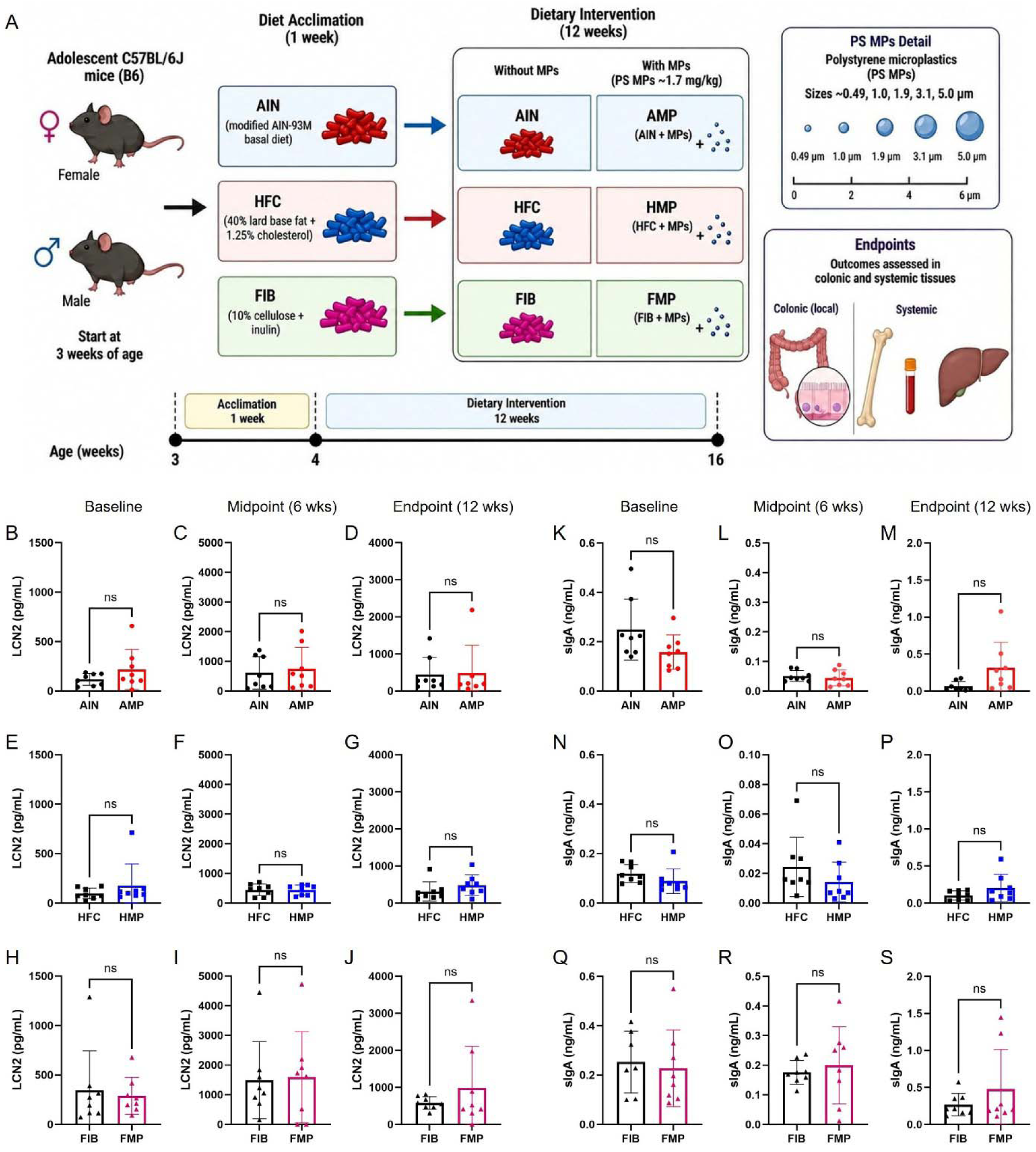
Dietary microplastic exposure does not induce overt intestinal inflammation. (A) Experimental design. Adolescent C57BL/6J female and male mice were acclimated at 3 weeks of age to one of three diets: modified AIN-93M basal diet (AIN), high-fat/high-cholesterol diet (HFC), or high-fiber diet (FIB). At 4 weeks of age, mice were maintained on these diets for 12 weeks with or without a physiologically relevant mixture of polystyrene microplastics (MPs; ∼0.5, 1.0, 2.0, 3.0, and 5.0 μm; total dose ∼1,7 mg kg⁻¹). Diets containing MPs are referred to as AMP (AIN + MPs), HMP (HFC + MPs), and FMP (FIB + MPs). (B-J), LCN-2 protein (pg/mL) measured across weeks −0 (B-D), −6 (E-G) and −12 (H-J) to evaluate intestinal inflammation in the stool of mice from each of the diets. (K-S), secretory IgA (sIgA) antibody (ng/mL) measured across weeks −0 (K-M), −6 (N-P) and −12 (Q-S) to evaluate intestinal inflammation in the stool of mice. Data are shown as mean ± s.d.; dots represent individual mice and analyzed using two-tailed unpaired Student’s t test. Statistical significance is indicated as ns, not significant.

Using dual-energy X-ray absorptiometry (DEXA), we observed significant diet- and sex-dependent effects of MP exposure on body composition. Inspection of fat mass revealed only male FMP mice had increased fat mass compared to male FIB mice (**Supplementary Fig. 2A-F**). For lean mass (**Supplementary Fig. 2G-L**), female AMP mice had an increase in lean mass compared to female AIN mice while male FMP mice displayed decreased lean mass compared to male FIB mice. Measuring whole-body bone mineral density (BMD) and composition (**Supplementary Fig. 2M-R**), DEXA only revealed significantly reduced BMD in AMP-fed females compared to AIN controls, whereas no significant differences were detected in the other groups. Beyond body composition, organ-specific effects of diet and MP exposure were also observed. We examined spleen mass and AMP female spleens were significantly larger that AIN controls; however, no differences in spleen mass were observed between the other dietary groups or sexes (**Supplemental Figure 3A-F**). Similarly, kidney weights were largely unaffected between the dietary groups or sexes (**Supplemental Figure 3G-L**). These findings suggest that MP exposure may adversely affect various tissues in a diet and sex-specific manner.

### Microplastics Alter Gut-Derived Endocrine Signaling Without Inducing Colonic Inflammation

Given past studies showing MP exposure leads to intestinal inflammation^17^, we next assessed colonic health in our dietary exposure model. Colon histopathology revealed uniformly low pathology scores across all diet and MP conditions (**Supplementary Figure 3M-R, Supplementary Table 3**). Most animals displayed intact intestinal architecture with no evidence of inflammation, and when changes were present, they were limited to mild submucosal edema (combined score = 1) without polymorphonuclear infiltration, goblet cell depletion, or epithelial injury. These minimal findings occurred sporadically across all groups and did not cluster by diet type, MP exposure, or sex. Additional assessment of fecal lipocalin-2 (LCN2) (**Figure 1B-J**) and secretory IgA (sIgA) (**Figure 1K-S**) at baseline (1 week after acclimation to diet without MPs), 6 weeks and 12 weeks revealed no significant changes between groups. Together, these findings indicate that chronic dietary MP exposure is not accompanied by overt colonic inflammation or epithelial injury.

Increased circulating inflammatory cytokines have also been linked to MP exposure including TNFα, MCP-1 and IL-6^51–57^. With the absence of intestinal pathology or overt inflammatory responses, we next asked whether MP exposure altered circulating inflammatory cytokines as well as gut-derived endocrine signals. Several studies have shown MP exposure increases TNFα levels^58–61^; however, TNFα did not differ between MP-exposed and control mice within any diet (**Fig. 2A-C**). Additionally, no change in MCP-1 was observed while IL-6 levels were only higher in HFC relative to HMP mice (**Supplementary Fig. 4A-F**). Interestingly, targeted plasma hormone profiling revealed a selective increase in circulating serotonin in AMP-fed mice compared with AIN controls (**Fig. 2D**). In contrast, serotonin concentrations did not differ between HFC and HMP groups or between FIB and FMP groups (**Fig. 2E,F**). Other hormones were largely unaffected by MP exposure: Glucose-dependent Insulinotropic Polypeptide (GIP) and resistin did not differ between MP-exposed mice and diet-matched controls across any dietary condition, while Glucagon-like Peptide-1 (GLP-1) was modestly reduced only in FMP compared with FIB mice (**Supplementary Fig. 4G-O**). The elevation of serotonin exclusively under AMP conditions suggests potential alterations in enterochromaffin cells, a major subset of EEC responsible for serotonin production.

**Figure 2.**
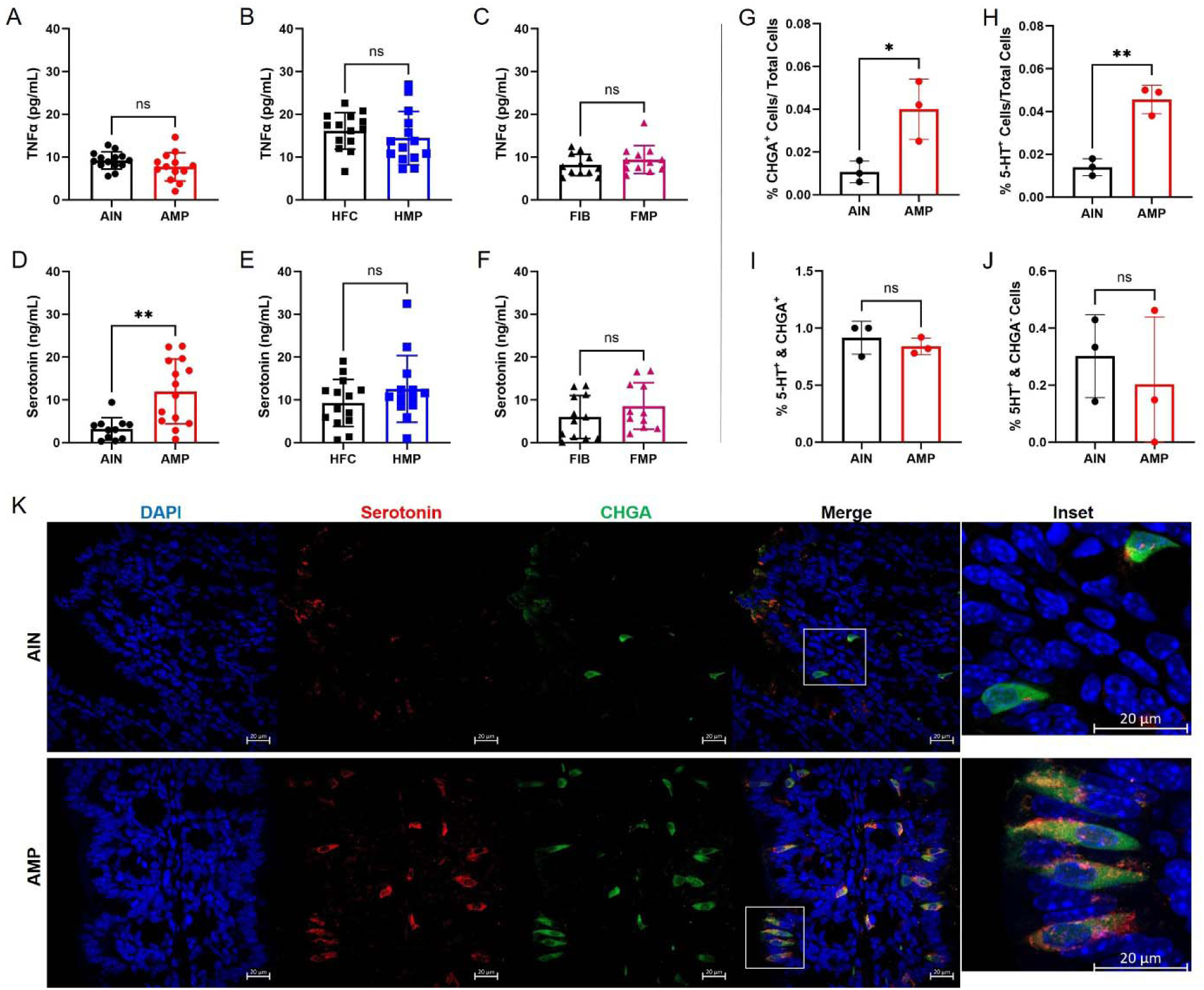
Dietary microplastic exposure enhances circulating serotonin and modulates enteroendocrine cell abundance. (A-C), Plasma TNFα levels in levels in AMP-fed mice compared with AIN controls (A), HFC and HMP (B) or FIB and FMP (C) groups. (D-F), Plasma serotonin (5-HT) AMP-fed mice compared with AIN controls (D), HFC and HMP (E) or FIB and FMP (F) groups. (G-J), Quantification of colonic enteroendocrine cells by immunofluorescence demonstrating increased proportions of CHGA^+^ cells (G) and 5-HT^+^ cells (H) in male AMP-fed mice relative to AIN male controls. Quantitative analysis showing no change in the fraction of serotonergic cells within the CHGA^+^5-HT^+^ population (I) and no increase in CHGA^−^/5-HT^+^ cells (J), indicating preserved enterochromaffin lineage identity despite increased enteroendocrine cell abundance. (K), Representative immunofluorescence images of proximal colon from AIN- and AMP-fed male mice stained for nuclei (DAPI, blue), serotonin (5-HT, red), and CHGA (green). Merged images and inset highlight increased CHGA^+^ and 5-HT^+^ cells in AMP-fed mice. Scale bars, 20 μm. Data are shown as mean ± s.d.; dots represent individual mice and analyzed using two-tailed unpaired Student’s t test. Statistical significance is indicated as *P < 0.05, **P < 0.01; ns, not significant.

### Dietary AMP Expands the Enteroendocrine Compartment and Enhances Serotonergic Output Without Lineage Skewing

Given the increase in circulating serotonin in the AMP group, we next examined EEC activity *in vivo*. Immunofluorescence staining of colonic tissue from AIN- and AMP-fed mice revealed increased chromogranin A (*CHGA)*^+^ and serotonin (5-HT)^+^ cells in male AMP-fed mice (**Fig. 2G-K**) as well as female AMP-fed mice (**Supplementary Fig. 5A,B**). Further quantitative analysis demonstrated that the fraction of serotonergic cells within the 5-HT^+^/*CHGA*^+^ *and the* 5-HT^+^/*CHGA*^-^ population were unchanged suggesting that MP exposure increases the abundance and/or detectability of EECs while preserving enterochromaffin lineage identity (**Fig. 2 I,J**). To further assess enterochromaffin cell-specific responses, we utilized *Piezo2*-GFP reporter mice^62^ where GFP is fused to the c-terminus of the endogenous Piezo2 protein and selectively labels a major subset of enterochromaffin cells^63^. AMP exposure increased 5-HT^+^ immunoreactivity per cell without altering the number or overall GFP intensity of Piezo2^+^ cells (**Supplementary Fig. 5C-E**), further suggesting enhanced serotonergic output within an otherwise stable enterochromaffin population.

To examine whether enhanced serotonergic signaling reflects intrinsic shifts in EEC differentiation, colonic organoids derived from AIN- and AMP-fed B6 mice were cultured and analyzed. Expression of enteroendocrine markers, including *Chga* (*i.e.*, chromogranin A) and the serotonin biosynthetic enzyme *Tph1* (*i.e.*, tryptophan hydroxylase 1), did not differ between AIN- and AMP-derived organoids in either stem cell-enriched or differentiated standard conditions (**Supplementary Fig. 6A-H**), indicating no sustained epithelial-intrinsic bias toward enteroendocrine lineage specification. These findings suggest proportional expansion or enhanced activity of the enteroendocrine compartment rather than selective skewing toward serotonergic differentiation.

To determine whether these effects are microbiota-mediated, we performed fecal microbiota transplantation (FMT)^36^ using stool collected at the endpoint from the AIN- and AMP-fed donor mice. At 12-weeks post-transfer, mice receiving AMP-derived microbiota displayed a significant increase of colonic CHGA^+^ cells relative to the AIN FMT group (**Supplementary Fig. 5F**), while 5-HT^+^ cells exhibited a non-significant upward trend (**Supplementary Fig. 5G**). These results implicate the microbiota shaped by the MP exposure is regulating EEC abundance, while suggesting that enhanced serotonergic signaling may require additional host or environmental inputs. Collectively, these findings support a model in which dietary MP exposure, in part through microbiota-dependent mechanisms, enhances enterochromaffin activity while expanding the overall enteroendocrine compartment without altering lineage identity.

### Single-nuclei RNA sequencing reveals a stable colonic epithelial composition without evidence of inflammation

To define epithelial-intrinsic mechanisms at single-cell resolution, we performed single-nuclei RNA sequencing (snRNA-seq) on colonic tissue from AIN- and AMP-fed mice. Unsupervised clustering resolved all major epithelial, immune, stromal, vascular, and neuromuscular compartments, with near-identical cell identities and UMAP structure across dietary conditions (**Fig. 3A; Supplementary Fig. 7A**), arguing against widespread transcriptional disruption. Epithelial cell composition was largely preserved, with no evidence for expansion of enterochromaffin cells (**Fig. 3B; Supplementary Fig. 7B**), consistent with organoid and reporter analyses. Enterochromaffin cells were robustly identified by expression of canonical markers (*Tph1*, *Chga*, *Tac1*), and differential abundance analysis indicated minimal changes across conditions. A modest reduction in goblet cells was observed under AMP conditions (**Supplementary Fig. 7B**), a change unlikely to account for increased serotonin given the absence of inflammatory gene signatures (**Supplementary Fig. 8A**) or histopathologic evidence of barrier disruption (**Supplementary Fig. 3M,N**). Given the use of snRNA-seq, transcript detection reflects nuclear RNA abundance, which may underrepresent cytoplasmic mRNAs with rapid turnover; however, relative expression differences between dietary groups remain interpretable at the level of transcriptional regulation^64–66^.

**Figure 3.**
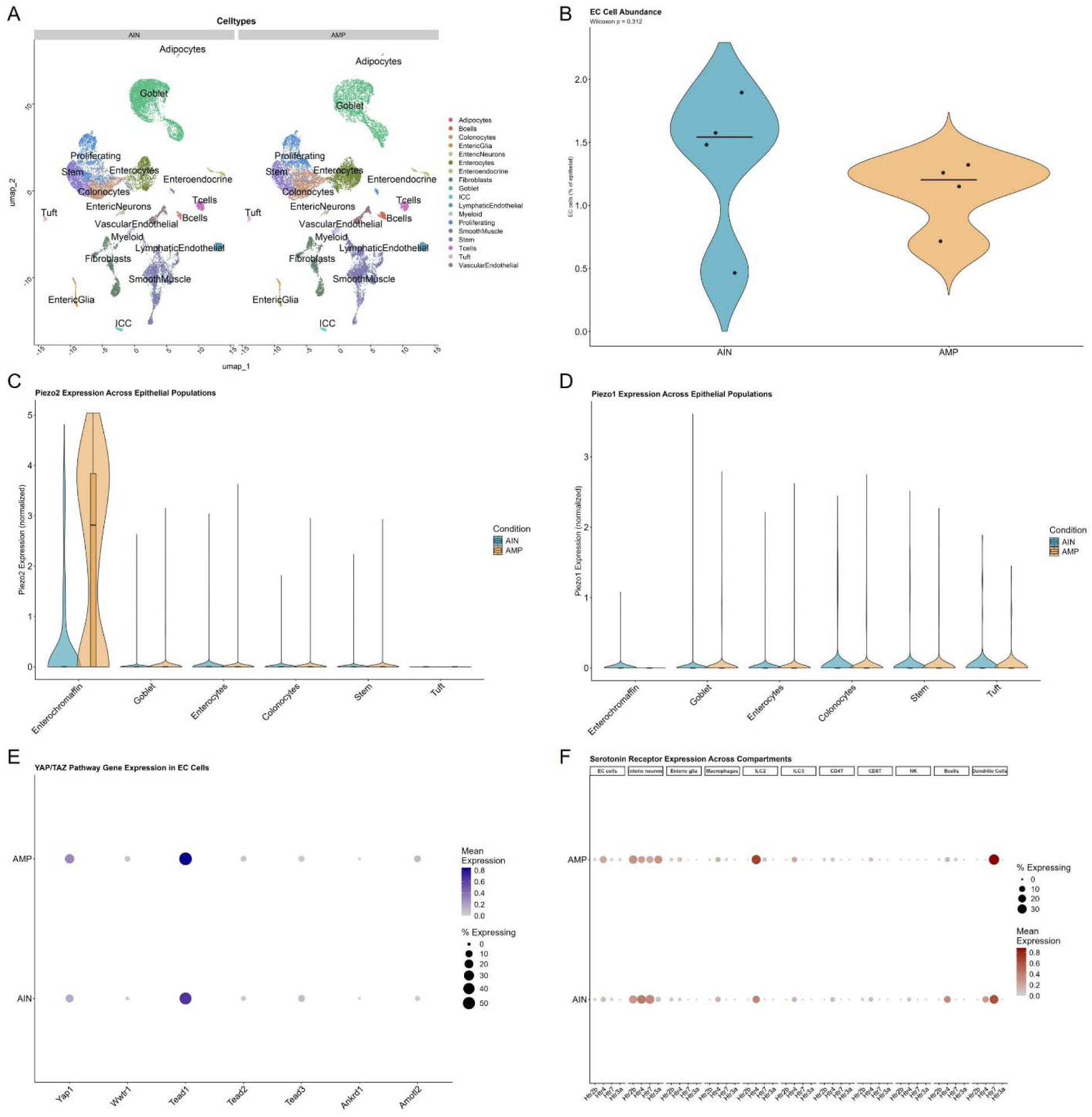
AMP exposure selectively modulates mechanosensory-associated transcripts in enterochromaffin cells without altering epithelial composition. (A), UMAP visualization of integrated single-nuclei RNA sequencing (snRNA-seq) data from colonic tissue of mice fed AIN or AMP diets (n = 4 biological replicates per group). Unsupervised clustering resolved major epithelial, immune, stromal, vascular, and neuromuscular compartments. Comparable cluster structure and cell identities between dietary conditions indicate preserved tissue architecture and minimal batch effects. (B), Relative abundance of enterochromaffin (EC) cells across diets. Violin plots show the percentage of EC cells per biological replicate. No significant difference was detected between AIN and AMP groups (Wilcoxon test). (C), *Piezo2* expression across epithelial populations. Violin plots show normalized single-cell expression stratified by cell type and diet. *Piezo2* transcripts were highly enriched in enterochromaffin cells and exhibited a rightward shift in AMP-fed mice, while remaining minimal in other epithelial populations. (D), *Piezo1* expression across epithelial populations. Piezo1 transcripts were rare (<1% of EC cells) and did not differ between diets. (E), Expression of YAP/TAZ pathway components in enterochromaffin cells. Dot plot showing mean expression (color intensity) and percentage of expressing cells (dot size) for *Yap1*, *Wwtr1* (Taz), *Tead1*, *Tead2*, *Tead3*, *Amot*, and *Amotl2*. Expression of core regulatory components was preserved across diets without coordinated induction of canonical YAP/TAZ target genes. (F), Serotonin receptor (Htr family) expression across colonic compartments. Dot plot depicts mean expression and percent-expressing cells across epithelial, neural, and immune populations. Receptor expression patterns were largely conserved between diets; H*tr3a* expression in enteric neurons showed a directional increase in AMP-fed mice, consistent with enhanced neuronal serotonergic responsiveness in a subset of animals. N=4 per group.

### snRNA-seq identifies mechanosensory-associated transcriptional changes in enterochromaffin cells

Serotonin release from enterochromaffin cells is mediated by the mechanosensitive ion channel Piezo2 that converts mechanical forces into the release of serotonin. We therefore set out to examine if enterochromaffin cells exhibited a focused transcriptional shift consistent with altered mechanosensory responsiveness. Interestingly, *Piezo2* transcripts were highly enriched in enterochromaffin cells and displayed a rightward shift in expression distribution under AMP conditions, while remaining minimal across other epithelial lineages (**Fig. 3C**); however, pseudo-bulk aggregation at the level of individual animals revealed substantial inter-animal variability, indicating that the magnitude of induction differed across biological replicates. The directionally consistent shift across animals, despite variable magnitude, suggests diet-responsive regulation with heterogeneous penetrance rather than stochastic noise. Notably, *Tph1* expression did not differ between diets (**Supplementary Fig. 7C**), indicating that increased serotonin production is unlikely to reflect transcriptional induction of the serotonergic biosynthetic machinery. We also examined the other mechanosensitive ion channel Piezo1 and found *Piezo1* expression was rare in enterochromaffin cells (<1%) and unchanged across other epithelial cell lineages between AIN and AMP diets (**Fig. 3D**). Together, these findings support a model in which dietary MP exposure enhances enterochromaffin secretory output through activity-dependent modulation of mechanosensory pathways, with proportional changes in EEC abundance or hormone content but without evidence of lineage reprogramming or induction of serotonergic gene expression programs.

Further analysis revealed enterochromaffin cells expressed the canonical transcriptional regulators *Yap1* and *Tead1* under both dietary conditions, consistent with a YAP/TAZ regulatory context capable of responding to mechanical cues (**Fig. 3E**); however, we did not observe coordinated induction of canonical YAP/TAZ target genes in AMP, suggesting engagement of mechanosensory signaling without widespread transcriptional reprogramming. Extending these observations, the mechanosensory-associated genes *Robo2* and its ligand *Slit3* exhibited increased expression across colonocytes, goblet cells, and enterochromaffin cells (**Supplementary Fig. 8B**), further supporting modulation of epithelial mechanosensory pathways in response to AMP exposure. Taken together, the data support modulation of mechanosensory responsiveness at the level of channel sensitivity or signaling efficiency rather than broad transcriptional remodeling of the enterochromaffin lineage.

Given the local increase in colonic 5-HT^+^ cells in AMP-fed mice, we next examined expression of serotonin receptor genes (Htr family) across epithelial, neural, and immune compartments to assess potential downstream responsiveness within the colonic microenvironment (**Fig. 3F**). Receptor expression patterns were largely preserved between diets; however, *Htr3a* transcripts in enteric neurons displayed a directionally consistent increase following AMP exposure. Pseudo-bulk aggregation at the level of individual animals revealed a similar fold change that did not reach statistical significance, reflecting inter-animal variability. These data suggest enhanced neuronal sensitivity to serotonergic input in a subset of animals. In addition to serotonergic receptor changes, differential abundance analysis revealed a significant reduction in cholinergic neurons in AMP-fed mice relative to controls (**Supplemental Fig. 7B**), indicating a selective decrease in this enteric neuronal subtype within the enteric neuronal network. Collectively, these findings support engagement of mechanosensory-associated pathways within enterochromaffin cells as a plausible epithelial mechanism linking dietary MP exposure to elevated peripheral serotonin, occurring in the absence of inflammation, epithelial restructuring, or widespread receptor reprogramming.

### Microplastic Exposure Promotes Hepatic Oxidative Stress Despite Minimal Effects of Steatosis

The liver also represents a central integrator of gut-derived signals via portal circulation and is a key component of the gut-liver axis through its regulation of nutrient, hormonal, and metabolic homeostasis. Among these gut-derived endocrine signals, peripheral serotonin is a potent regulator of hepatic lipid metabolism, mitochondrial activity, and inflammatory signaling ^67–70^. Consistent with our observation of increased EEC density and elevated colonic and circulating serotonin, hepatic serotonin was also increased in AMP livers compared to AIN controls (**Supplementary Fig. 5H**). Interestingly, the liver has been identified to be a site of MP accumulation, with reports suggesting links to steatosis and inflammatory responses, as recently reviewed^71^. Histological assessment of hepatic steatosis revealed minimal effects of MP exposure under normal- or fiber-based diets: AIN, FIB, and FMP livers showed no steatosis, and only one of five AMP mice exhibited mild (grade 1) steatosis (**Fig. 4A-F; Supplemental Table 4**). In contrast, high fat/high cholesterol diets produced pronounced sex-dependent pathology. HFC and HMP females exhibited only mild (grade 1) steatosis, whereas males in both groups consistently showed severe (grade 3) steatosis, with no apparent MP-specific exacerbation beyond that induced by the high fat/high cholesterol diet itself (**Supplementary Table 4**). No differences in liver mass were observed between dietary groups or sexes (**Fig. 4G-L**). Because MPs were administered within the dietary formulations rather than by OGG, differences in hepatic MP burden or distribution could contribute to the absence of steatosis observed in our model. Therefore, we next sought to visualize MP-associated signals in paraffin-embedded liver tissue sections using a *conjugated polymer nanoparticle-based dye, previously validated for stable, long-term microplastic detection*^72^. Using this approach, polymer-positive signals were readily detected in MP-exposed animals (**Fig. 4M-R**), confirming hepatic accumulation of MPs in mice in all dietary formulations containing MPs.

**Figure 4.**
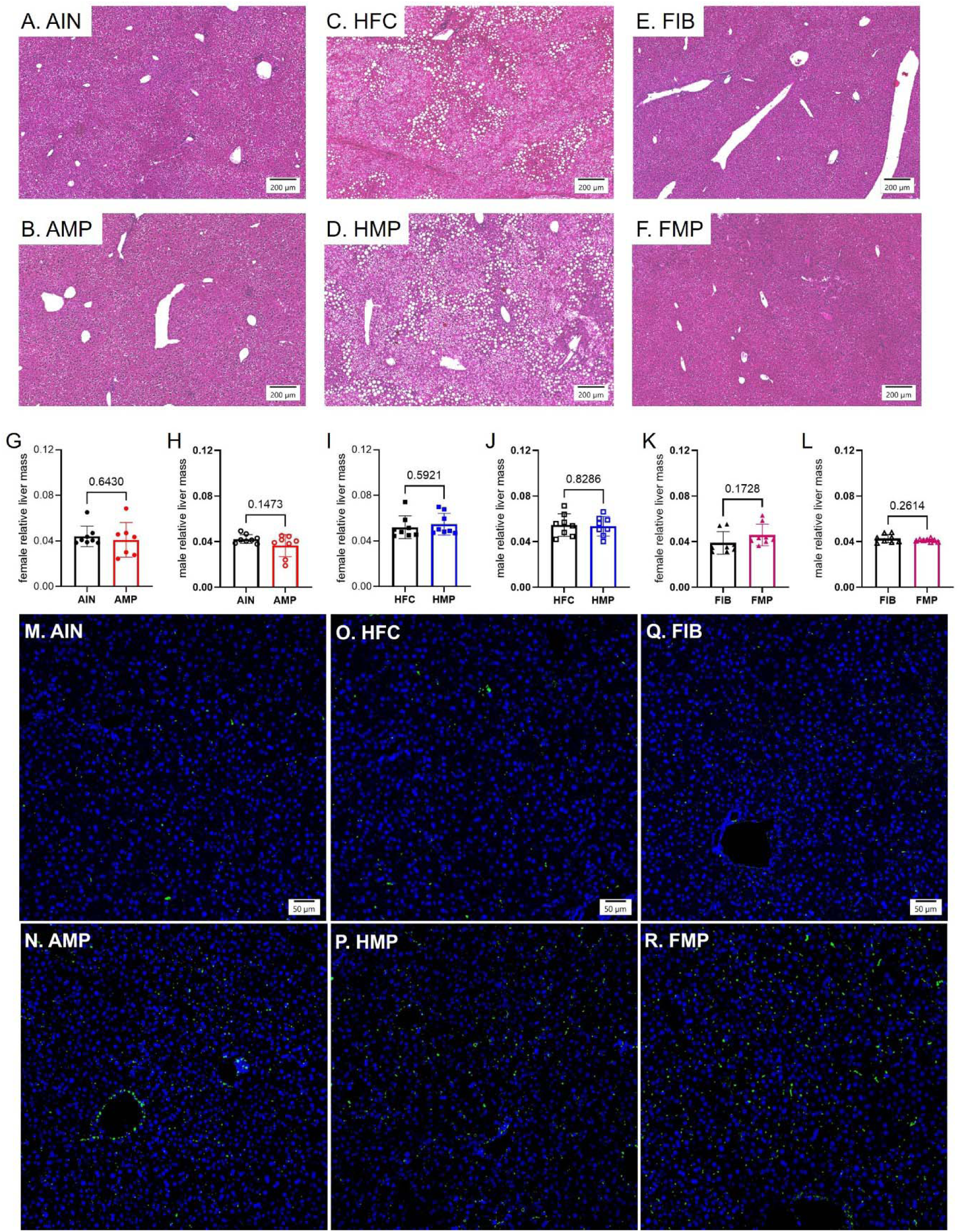
Hepatic histology, liver mass, and microplastic-associated signals in mice fed experimental diets with or without microplastics. (A-F) Representative H&E-stained liver sections from mice fed AIN (A), AMP (B), HFC (C), HMP (D), FIB (E), or FMP (F) diet. Liver sections were evaluated using the NASH Clinical Research Network Scoring System (scores reported in Supplementary Table 4). (G-L) Relative liver mass in female (G,I,K) and male (H,J,L) mice from the indicated dietary groups. Each symbol represents an individual animal. (M-R) Representative fluorescence images of paraffin-embedded liver sections stained with a conjugated polymer nanoparticle-based dye for detection of polymer-associated signals. Liver sections from mice fed AIN (M), AMP (N), HFC (O), HMP (P), FIB (Q), or FMP (R) diet. Scale bars, 50 μm. Data are shown as mean ± s.d.; dots represent individual mice and analyzed using two-tailed unpaired Student’s t test.

Since histological steatosis was only modestly altered by MP exposure but MP were present, we next examined markers of hepatic oxidative stress and inflammation. Liver sections were stained for 4-hydroxynonenal (4-HNE), a marker of lipid peroxidation and oxidative stress, and myeloperoxidase (MPO1). Under normal dietary conditions, AMP livers exhibited visibly increased 4-HNE immunoreactivity compared to AIN controls, despite minimal histologic evidence of steatosis (**Fig. 5A-B**). Similarly, FMP livers demonstrated increased diffuse 4-HNE staining relative to FIB controls (**Supplemental Fig. 9**). In contrast, MPO1 staining showed comparatively modest differences between groups across dietary conditions, suggesting limited inflammatory cell infiltration (**Supplemental Fig. 9**). As expected, HFC and HMP liver sections exhibited pronounced steatotic morphology and elevated baseline oxidative stress with only modest increase in 4-HNE staining observed in HMP tissues relative to HFC controls (**Supplemental Fig. 9**). Collectively, these findings indicate that dietary MP exposure consistently promotes hepatic oxidative stress across nutritionally distinct dietary conditions. Nevertheless, there was comparatively limited changes in overt inflammatory staining or steatosis severity. Take together, this suggests chronic MP exposure may alter hepatic physiology through coordinated serotonergic and oxidative stress-associated pathways.

**Figure 5.**
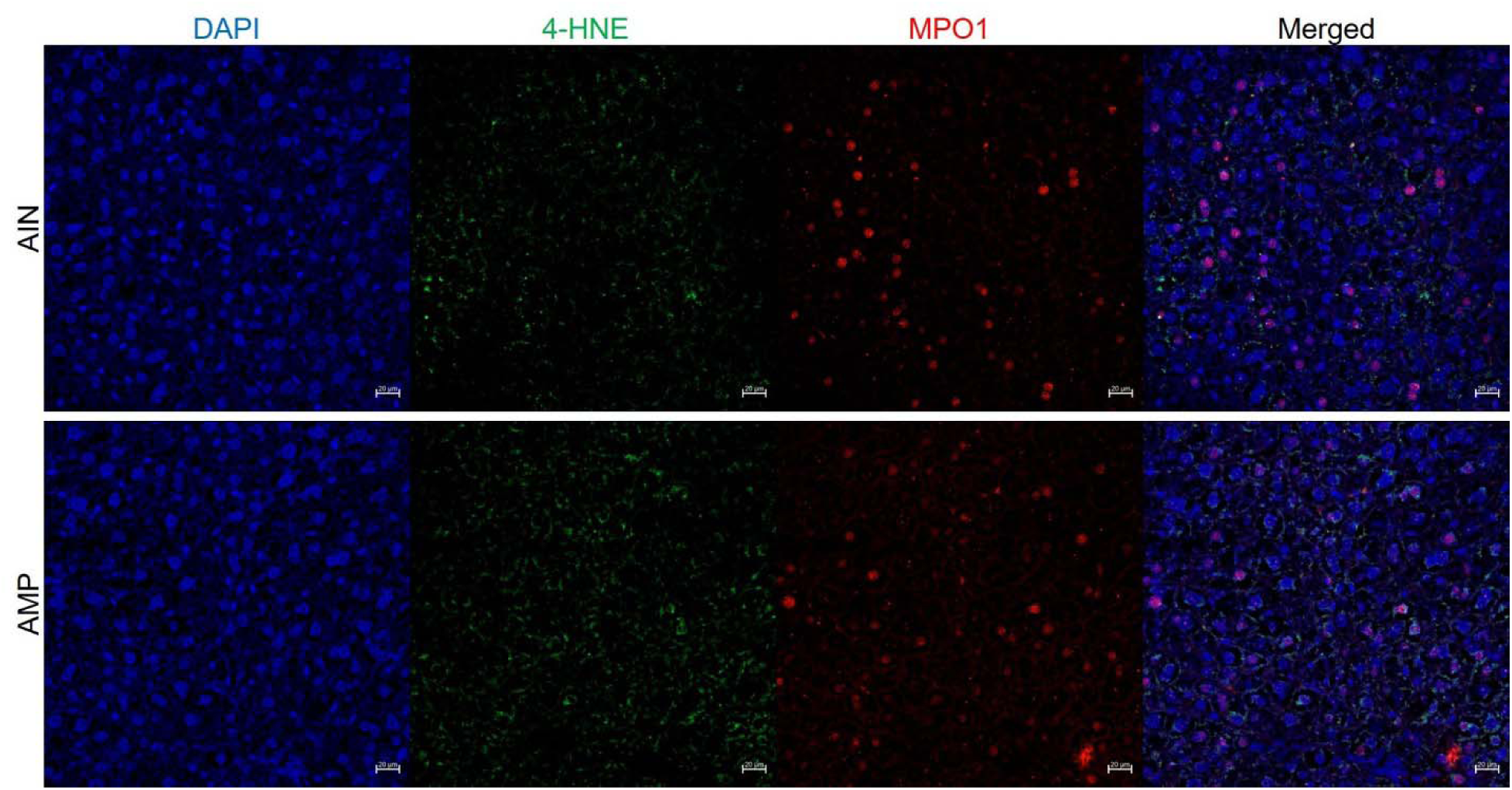
Representative immunofluorescence detection of hepatic oxidative stress in AIN and AMP mice. (A-B) Representative liver sections from AIN (A) and AMP (B) mice stained for 4-hydroxynonenal (4-HNE; green) and myeloperoxidase (MPO1; red), and nuclei counterstained with DAPI (blue). Merged images are shown in the far-right panels. Scale bars, 20 μm.

### Chronic Dietary MP Exposure Disrupts Cortical and Trabecular Bone Architecture in a Sex- and Diet-Dependent Manner

Given the increase in colonic and peripheral serotonin and its association with bone loss as well as the limited resolution of DEXA for distinguishing cortical from trabecular compartments, we next conducted high-resolution micro-computed tomography (µCT) of the femur and vertebrae. This allowed us to localize and characterize MP-related skeletal changes. µCT analysis demonstrated that chronic MP exposure selectively degraded trabecular microarchitecture in both sexes, with additional cortical deficits in females. In AMP-fed females, cortical area and total cross-sectional area were reduced without changes in cortical thickness, this was accompanied by lower trabecular BV/TV and trabecular number, greater trabecular separation, and reduced connectivity (**Fig. 6A-E, Supplementary Fig. 10A-C**). AMP-fed males also displayed substantial trabecular deterioration including reduced BV/TV, trabecular thickness, and trabecular number, with increased separation and diminished connectivity density; however, cortical structure remained unchanged (**Fig. 6F-J, Supplementary Fig. 10D-F**). To identify potential mechanisms, we profiled genes regulating bone formation (*Runx2*, *Alpl*), Wnt signaling (*Dkk1*), and inflammation (*Il1a*). In AMP females, combined cortical and trabecular bone loss was associated with an upregulation of *Dkk1* and *Il1a* (**Fig. 6K-N**), consistent with Wnt signaling inhibition and inflammation-driven osteoclast activation, suggesting enhanced bone resorption and suppressed bone formation. Notably, expression of *Runx2* and *Alpl* was unchanged in this group (**Fig. 6K-N**). In AMP males, where trabecular bone loss occurred without cortical involvement, *Runx2*, a key driver of osteoblast differentiation, was reduced (**Fig. 6O-R**), suggesting impaired bone formation is the dominant mechanism, while *Alpl*, *Dkk1* and *Il1a* genes remained unchanged (**Fig. 6O-R**). Spinal µCT revealed more modest, diet-dependent changes. AMP-fed females showed only reduced trabecular thickness (**Supplementary Fig. 10G-K**), whereas AMP-fed males exhibited increased trabecular thickness coupled with reduced connectivity density (**Supplementary Fig. 10L-P**).

**Figure 6.**
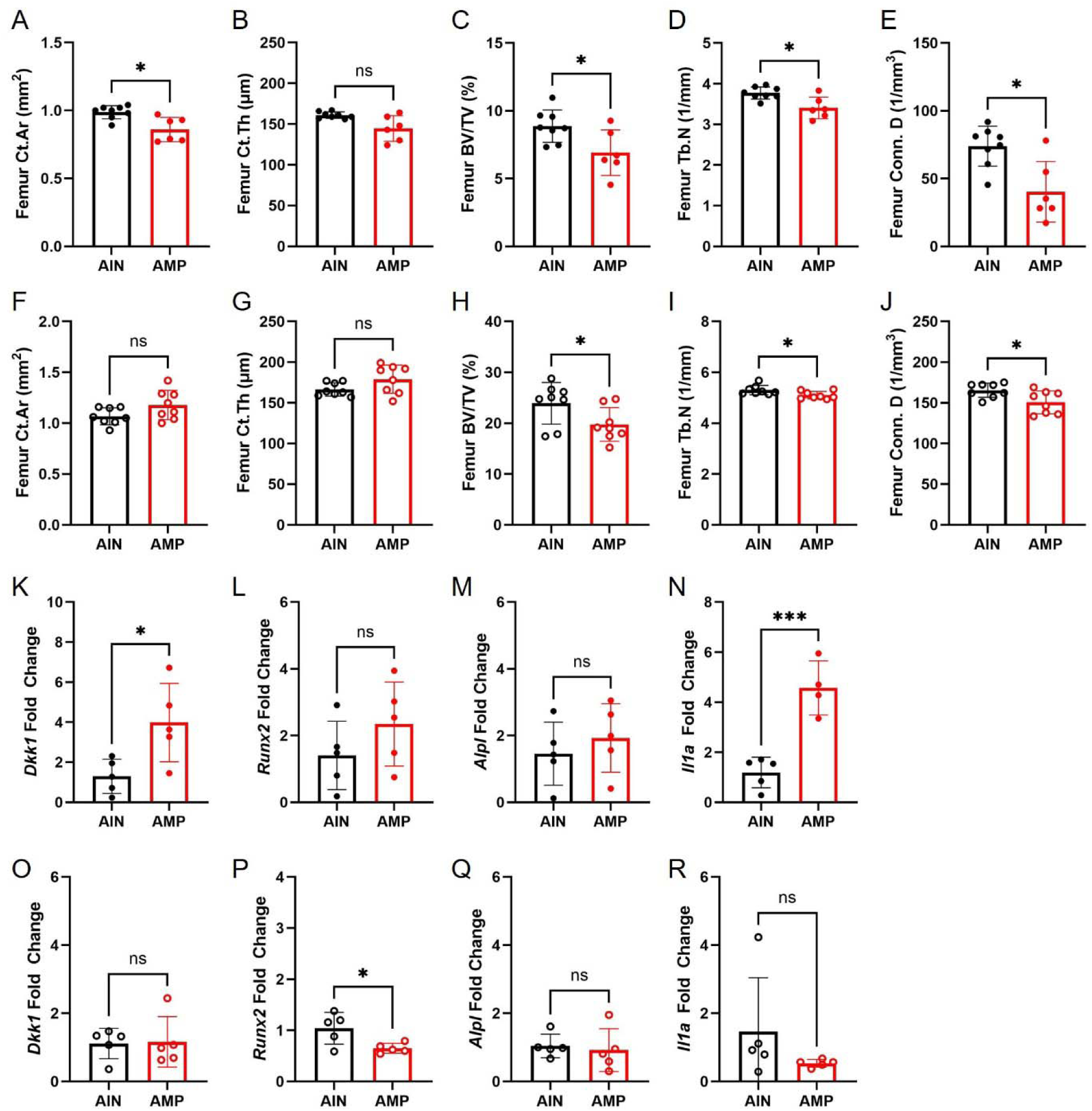
Dietary microplastic exposure disrupts bone microarchitecture in a sex-specific manner. (A-E), Femoral µCT analysis in females (closed circles) showing reduced cortical area (Ct.Ar) and cortical thickness (Ct.Th) (A,B), together with decreased trabecular bone volume fraction (BV/TV) (C), trabecular number (Tb.N) (D), and connectivity density (Conn. D) (E) in AMP-fed mice relative to AIN controls. (F-J), Femoral µCT analysis in males (open circles) demonstrating preserved cortical parameters (F,G) but reduced trabecular BV/TV (H), Tb.N (I), and Conn. D (J) in AMP-fed mice. (K-N), Relative mRNA expression of *Dkk1* (K), *Runx2* (L), *Alpl* (M), and *Il1a* (N) in femoral bone from females. (O-R), Relative mRNA expression of *Dkk1* (O), *Runx2* (P), *Alpl* (Q), and *Il1a* (R) in femoral bone from males. Data are mean ± s.d.; dots represent individual mice. Statistical significance is indicated as \**P* < 0.05, \*\*\**P* < 0.001; ns, not significant.

In contrast, MP exposure under high-fat or high-fiber diets produced more limited femoral effects: HMP females showed no MP-related differences in cortical or trabecular structure compared to HFC females (**Supplementary Fig. 11A-H**), whereas HMP males exhibited reduced trabecular BV/TV and trabecular thickness with preserved cortical morphology (**Supplementary Fig. 11I-P**). HMP females showed no changes in *Dkk1*, *Runx2*, *Alpl*, or *Il1a* (**Supplementary Fig. 12A-D**), consistent with minimal skeletal changes. In contrast, HMP males exhibited reduced *Runx2* and *Alpl*, indicating impaired osteoblast differentiation and maturation, while markers of Wnt signaling (*Dkk1)* and inflammation (*Il1a)* remained unchanged (**Supplementary Fig**. **12E-H**). For the high fiber groups, FMP females also showed no detectable impact of MP exposure (**Supplementary Fig**. **13A-H**), while FMP-fed males displayed coordinated deficits across both compartments, including reduced cortical area, total area, and cortical thickness, together with lower trabecular BV/TV and trabecular thickness (**Supplementary Fig. 13I-P**). FMP-fed females and males had no significant changes in *Alpl*, *Runx2*, or *Dkk1* expression while *Il1a* was undetectable in both FIB and FMP samples (**Supplementary Fig. 14A-F**). These findings suggest that cortical and trabecular bone loss observed in FMP-fed males may be driven by alternative, as yet undefined mechanisms. No spine parameters were altered in HMP-females (**Supplementary Fig. 11Q-U**) or in FMP-fed females (**Supplementary Fig. 13Q-U**), while HMP-fed males displayed reduced BV/TV and trabecular thickness (**Supplementary Fig. 11V-Z**) and FMP-fed males showed decreased trabecular thickness with increased connectivity density (**Supplementary Fig. 13V-Z**). Collectively, these data demonstrate that chronic MP ingestion disrupts bone homeostasis in a sex- and diet-specific manner, with the trabecular compartment most consistently affected. Notably, the selective elevation of serotonin, observed only under AMP conditions, coincided with the most pronounced bone loss observed in AMP-fed animals, supporting a potential gut endocrine contribution to MP-associated skeletal deficits.

### Evidence of Microplastic Accumulation in Human Bone and Functional Impairment of Osteoblast Mineralization

*O*rthopedic prostheses and implants have been shown to generate plastic debris at the bone-implant interface and are reported to affect bone biology^73^. Recent reports using *microphotography and Raman spectroscopy* have identified MP in human skeletal tissues^34^. Given our results showing MP exposure caused skeletal defects, we sought to determine whether MPs are detectable in human skeletal tissue in individuals without prostheses. Thus, we analyzed histological sections of the human distal femur (from a commercial source, donor information in **Supplementary Table 6**). W*e applied the above mentioned conjugated polymer nanoparticle-based dye*^72^, *to visualize MP-positive signals in intact human distal femur sections.* MP-positive signals were detected in distal femurs from five independent human donors (**Figure 7A-C; Supplementary Fig. 15**). The specificity of this staining approach was supported by parallel analysis of liver tissue from mice maintained on control diets or MP-supplemented diets, in which polymer-positive signals were readily detected in MP-exposed animals (**Fig. 4 M-R**). The detection of polymer signals in mineralized human tissue suggests that MPs can persist and accumulate in the skeleton, raising the possibility that environmental MP exposure may be an unrecognized risk factor for bone fragility.

**Figure 7.**
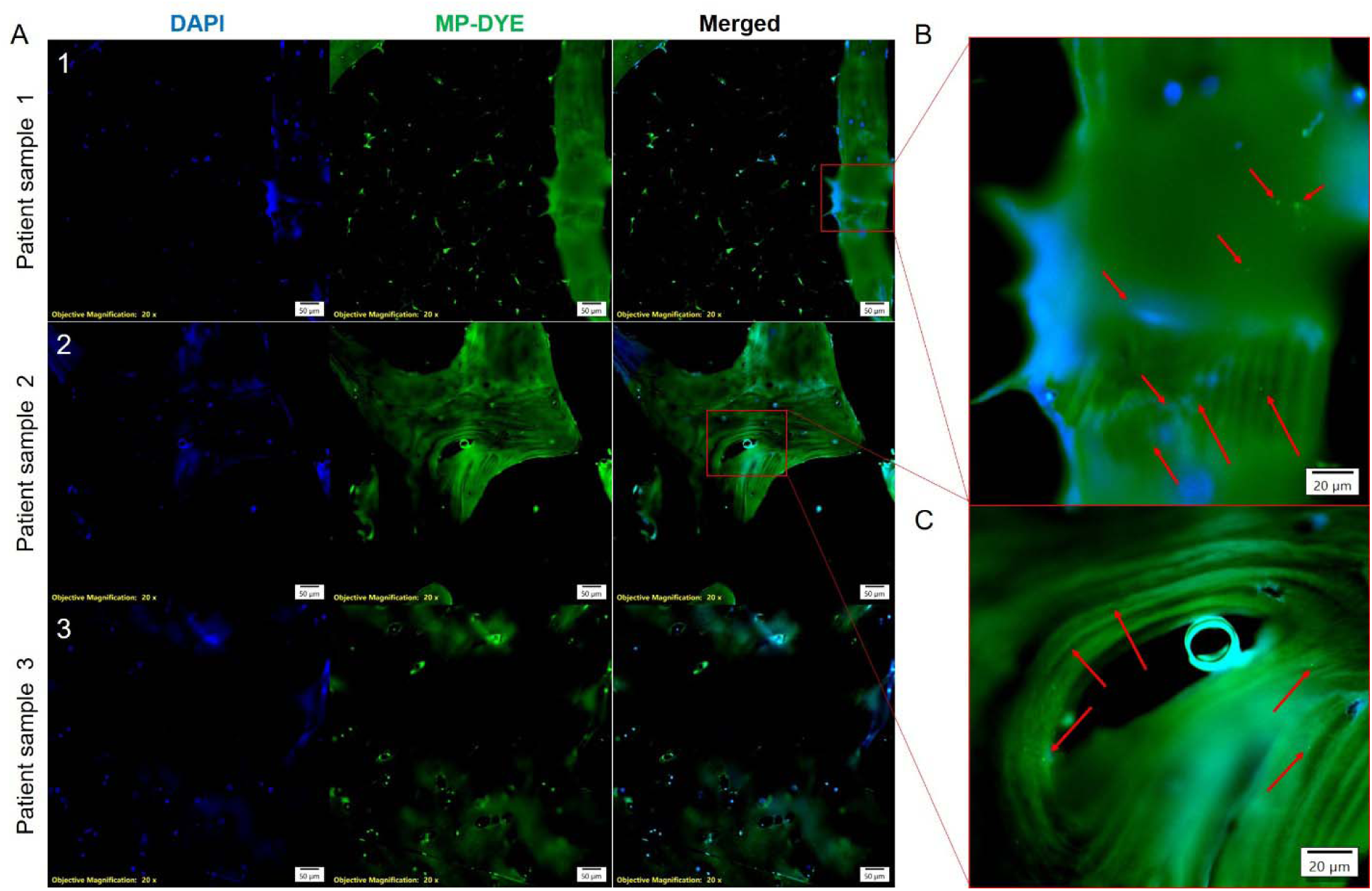
Polymer detection in human distal femur tissue from donors without orthopedic prostheses. (A) Representative fluorescence images of histological sections from distal femurs from three of the independent human donors (Patients 1-3; donor information in Supplemental Table 6) stained with DAPI (blue) to label nuclei and a conjugated polymer nanoparticle-based MP dye (MP-DYE, green). Merged images demonstrate discrete MP-positive signals within intact bone tissue. Objective magnification: 20X. Scale bars, 50 µm. (B,C), Higher-magnification views of boxed regions in (A) highlighting (red arrows) irregular MP-positive signals (green) embedded within the bone matrix. Scale bars, 20 µm.

Consistent with this possibility, *in vitro* studies have demonstrated that PS MPs impair osteogenic differentiation and induce senescence in MC3T3-E1 pre-osteoblasts^30^. To examine the direct effects of MPs and serotonin on human bone-forming cells, primary human osteoblasts from three independent donors (donor information in **Supplementary Table 8**) were cultured for 9 days in the presence of 1- or 5-µm PS MP (1 µg mL^-1^, **Supplementary Table 2**) or serotonin (10 or 50 ng mL^-1^), followed by washing and induction of mineralization (**Supplementary Fig. 16A**). In osteoblasts derived from two donors in their mid-70s, transient exposure to PS MPs resulted in a marked and sustained reduction in calcium deposition compared with untreated, donor-matched controls (**Supplementary Fig. 16B**). In these same donors, serotonin exposure also impaired mineralization, although sensitivity varied, with suppression observed at 50 ng mL^-1^ in one donor and 10 ng mL^-1^ in the other. In contrast, osteoblasts from a third donor in their mid-60s exhibited preserved mineralization following MP or serotonin exposure, indicating inter-individual variability in osteoblast sensitivity to serotonergic and microplastic perturbation. Together, these findings demonstrate that MPs can persist in human bone tissue and disrupt osteoblast mineralization capacity in a donor-dependent manner.

## DISCUSSION

In this study, we investigated how chronic dietary MP exposure influences gut-endocrine signaling and skeletal homeostasis across physiologically relevant dietary contexts. In the standard base diet (AIN and AMP), we demonstrate the MP ingestion leads to a consistent increase EECs and an enterochromaffin-derived serotonergic output in the colon, accompanied by enhanced mechanosensory signaling within enterochromaffin cells including modulation of Piezo2-associated pathways. These changes occurred in the absence of epithelial lineage reprogramming, overt inflammation, or broad transcriptional remodeling. This points to a functional rather than developmental shift in EEC activity. Additionally, AMP-shaped microbiota was partially responsible for this phenotype by enhancing EECs. We also found MPs accumulate in the liver regardless of dietary formulation and AMP conditions have increased levels of hepatic serotonin; however, MP-induced steatosis was not observed in our 12-week model of exposure. Other systemic consequences of MP exposure were sex- and diet-dependent showing altered body composition and selective reduction in BMD and trabecular microarchitecture, most prominently under normal dietary conditions in females. Mechanistically, skeletal alterations were associated with distinct molecular signatures indicative of altered Wnt signaling, inflammation, or impaired osteoblast differentiation depending on sex and dietary conditions. Finally, we identify the presence of MP-associated signals in human distal femur tissues and demonstrate that transient exposure of MPs impairs osteoblast mineralization in a donor-dependent manner. Collectively, these findings extend the concept of dietary constituents beyond nutrients and bioactive metabolites to include synthetic particulate materials capable of modulating gut endocrine and metabolic physiology.

MPs are reported to be endocrine disruptors in the adrenal gland, ovaries and thyroid ^19–24^, our data extends this to the GI tract. We found AMP exposure increased serotonergic immunoreactivity; however, we did not observe increase in Piezo2-GFP reporter intensity or Piezo2^+^ cell number. If AMP exposure increased total Piezo2 abundance, a proportional increased in GFP reporter signal would be expected. However, anti-GFP immunofluorescence revealed no overt change in reporter intensity with enterochromaffin cells under AMP conditions. This distinction is notable because the Piezo2-GFP reporter reflects total fused protein abundance rather than channel localization, membrane insertion, or mechanosensitive activity. Thus, enhanced serotonergic output may arise through post-translational regulation of Piezo2, including altered membrane residency, trafficking, gating sensitivity, or protein stabilization, without detectable changes in total GFP signal^74–76^. Alternatively, AMP exposure may preferentially augment serotonin synthesis or release of Piezo2 activation rather than expanding the Piezo2^+^ enterochromaffin population itself. These findings raise the possibility that MP exposure alters enterochromaffin cell state or excitability rather than lineage allocation.

Dietary contexts also appeared to strongly influence the serotonergic response to MP exposure. Notably, HMP diet failed to further increase serotonin levels in mice consuming MPs in the presence of a high-fat/high-cholesterol diet. A high-fat diet has been shown to silence EEC through microbiota-dependent mechanisms^77^. Another potential explanation is that lipid-rich diets may suppress mechanosensory signaling through altered Piezo channel function. Prior studies have demonstrated that dietary fatty acids can enhance Piezo2 channel inactivation and reduce mechanically evoked signaling, indicating that lipid environments can substantially influence Piezo2-dependent mechanotransduction^78–80^. In contrast, the lack of an additive serotonergic effect in the high-fiber diets (*e.g.*, FIB and FMP) may reflect a ceiling effect in enterochromaffin stimulation. Fermentable fiber is a potent modulator of gut microbial metabolism that results in increased production of short-chain fatty acids and microbial metabolites that enhance enterochromaffin signaling and serotonin biosynthesis through regulation of TPH1 expression and enterochromaffin cell activity^81, 82^. Under these conditions, the enterochromaffin compartment may already exist in a relatively activated state, which may limit the capacity of additional MP-associated mechanical stimulation to further elevate serotonin production.

The FMT studies further suggest that microbiota-dependent and mechanical components of MP exposure contribute distinct but complementary signals. While transfer of AMP-associated microbiota increased CHGA^+^ EECs, the increase in 5-HT^+^ cells did not reach significance, indicating that microbiota remodeling alone may be insufficient to fully recapitulate the serotonergic phenotype observed during direct AMP exposure. One possible model is that MP-shaped microbial communities promote expansion or priming of the enteroendocrine compartment, while the persistent physical presence of MPs within the intestinal lumen provides mechanical stimulation that enhances enterochromaffin cell activity and serotonergic output. Prior work from Beyder and colleagues demonstrated that intraluminal microbeads can directly stimulate Piezo2-dependent mechanosensory pathways and alter GI motility^63, 83^. Although these microbeads were ∼100-microns, this supports the possibility that MPs act in a similarly function as a chronic luminal mechanical stimulus within the gut environment. This mechanism would be consistent with the established mechanosensory role of Piezo2 in enterochromaffin cells and may explain why microbiota transfer alone incompletely phenocopied the serotonin changes induced by AMP exposure. An additional consideration is that residual MPs have remained within the donor fecal preparations used for FMT. Although fecal pellets were resuspended in sterile PBS and subjected to gentle centrifugation prior to transfer, only the supernatant fraction was collected for gavage, raising the possibility that smaller suspended MP particles persisted within the inoculum. Given that the recipient mice were analyzed 12-weeks after transplantation, it is unlikely that transient carryover alone fully accounts for the observed phenotype. Also, the relatively rapid turnover dynamics of intestinal epithelial and EEC populations, which are generally renewed over days to weeks, and the persistence of increased CHGA^+^ cell abundance 12 weeks after transplantation suggests a microbiota-dependent remodeling^84^.

Additional snRNA-seq analysis identified a marked reduction in goblet cells and cholinergic enteric neurons following MP exposure. Goblet cells regulate mucus rheology and luminal biomechanics, whereas cholinergic enteric neurons regulate mucus secretion and epithelial sensory signaling. These findings suggest that MP exposure disrupts the mucus-neuroepithelial interface that normally buffers and coordinates responses to luminal mechanical stimuli. Notably, these cellular changes occurred in the absence of overt colonic inflammation, indicating a form of non-inflammatory tissue remodeling. Given the reciprocal interactions among these cells, loss of goblet cells and cholinergic enteric neuronal signaling may alter the biomechanical and sensory environment of the colonic mucosa, thereby enhancing EEC responsiveness to mechanical stimuli. Taken together, these findings support a dual-hit model whereby MP exposure promotes microbiota-dependent remodeling of the enteroendocrine compartment while simultaneously providing chronic luminal mechanical stimulation. We envision that these combined effects alter the biomechanical and microbial environment of the colon, resulting in heightened enterochromaffin cell activity and serotonergic output through Piezo2-dependent mechanosensory pathways.

Beyond its local effects on intestinal physiology, enterochromaffin-derived serotonin functions as a major endocrine signal capable of influencing peripheral organ systems including the liver and skeletal system^26, 67, 85–87^. Consistent with this concept, we observed increased hepatic serotonin levels in AMP-exposed mice despite the absence of overt steatosis as well as skeletal defects in both the femur and vertebrae. Serotonin has been implicated in regulation of hepatic metabolism, regeneration and fibrosis through a gut-liver axis^87^. Gut-derived serotonin has also been reported to suppress osteoblast function and bone formation^86^. Although our study was not designed to establish causality with regards to bone loss, the concurrence of enhanced enterochromaffin serotonergic output, elevated peripheral and hepatic serotonin, and impaired skeletal parameters raises the possibility that chronic MP exposure engages a systemic gut-derived serotonergic axis that contributes to metabolic and skeletal adaptations.

As for the impact of plastics on bone, synthetic polymers are extensively used in orthopedic prostheses and implants, where long-term mechanical wear inevitably generates plastic debris at the bone-implant interface. Indeed, ultra-high molecular weight polyethylene (UHMWPE), a common material in joint replacements, has been shown to produce wear particles. These particles induced a catabolic, bone-resorbing phenotype in osteocytes without causing osteocyte death by directly introducing UHMWPE particles into the surgical site, although the particle size was unreported^73^. While plastic wear particles from prosthetic materials are known to affect bone biology, more recent studies have been investigating the effects of MPs under non-iatrogenic conditions. These studies implicate MP as inducers to systemic inflammation and bone marrow toxicity that ultimately contribute to skeletal dysfunction^25, 30–34^; however, the route of MP exposure may critically influence the observed outcomes. Interestingly, we could detect MP-associated signals in human bones further supporting previous work^34^. Although our study does not establish causality in humans, it identifies a gut-bone endocrine axis responsive to dietary MPs and highlights mechanosensory enterochromaffin signaling as a potential interface between environmental materials and host physiology. Whether similar MP-induced serotonergic responses occur in humans and how dietary composition modulates skeletal vulnerability or GI motility warrant further investigation. Nevertheless, these findings support a model in which dietary MPs act as modulators of enteroendocrine function with downstream, context-dependent effects on hepatic and skeletal homeostasis.

## STUDY LIMITATIONS

Future studies employing pharmacologic or genetic modulation of peripheral serotonergic signaling will be required to determine whether gut-derived serotonin causally mediates the skeletal consequences of chronic MP exposure. Likewise, although the observed enterochromaffin phenotype is consistent with altered mechanosensory signaling, direct interrogation of Piezo2 function was not performed and therefore the contribution of Piezo2-dependent pathways remains inferential. Classical systemic markers of bone turnover such as P1NP, osteocalcin, and collagen type I C-telopeptide (CTx) were not measured, and dynamic histomorphometry was not performed. Nevertheless, the transcriptional changes observed in bone tissue are consistent with MP exposure influencing both osteoblast and osteoclast regulatory pathways. Notably, expression of Tnfs11 (RANKL) in bone was unchanged across AMP and HMP dietary conditions (data not shown), suggesting that the skeletal phenotypes observed here are unlikely to be explained solely by alterations in canonical osteoclastogenic signaling. Instead, increased expression of inflammatory mediators and Wnt signaling antagonists in AMP females is consistent with enhanced bone resorption, whereas reduced Runx2 expression in AMP males suggests impaired osteoblast differentiation. Future studies integrating serum bone turnover markers, dynamic histomorphometric analyses, and mechanistic manipulation of serotonergic and mechanosensory pathways will be required to define how MP exposure alters bone remodeling dynamics.

## Acknowledgments

We would like to acknowledge the UNM Comprehensive Cancer Center Support Grant NCI P30CA118100 and the HSC Shared Equipment for use of the Olympus iX83 Yokogawa Spinning Disk Confocal Microscope and Evident APEXVIEW APX100 Benchtop Fluorescence Microscope within the Advanced Light Microscopy Resource, Analytical and Translational Genomics (ATG) core; Bioinformatics and Data Science core; and Human Tissue Repository and Tissue Analysis core. We thank Michael L. Paffett at the UNM Advanced Light Microscopy Resource for training and guidance with confocal imaging. During manuscript preparation, an AI-based language tool was used to improve readability and clarity of text. All content was reviewed and approved by the authors, who take full responsibility for the final manuscript. Graphical figures were created with BioRender.com.

## Author contributions

Conception and design: ASR, SP, EFC

Development of methodology: ASR, SP, RP, EFC

Acquisition of data: ASR, SP, SP, HYD, JAR, OMO, BBM, SSG, CME, FTL, JO, CC, RL, KC, AC

Analysis and interpretation of data: ASR, SP, OMO, RL, JMG, AC, RP, EFC

Writing, review, and/or Revision: ASR, SP, RP, EFC

Administrative, technical, or material support (i.e., reporting or organizing data, constructing database): ASR, SP, OMO, JGI, MC, KC, RL, JGC, SL, RP, EFC

Study supervision and funding acquisition: EFC

## Data availability statement

All primary data associated with this study are present in the paper, the Supplementary Materials or will be deposited upon publication.

