## Supplemental Figures for "Dietary Microplastics Engage Gut Mechanosensory-Endocrine Signaling and Disrupt Bone Homeostasis"

Aaron S. Romero et al.,

Supplemental Figures 1-16

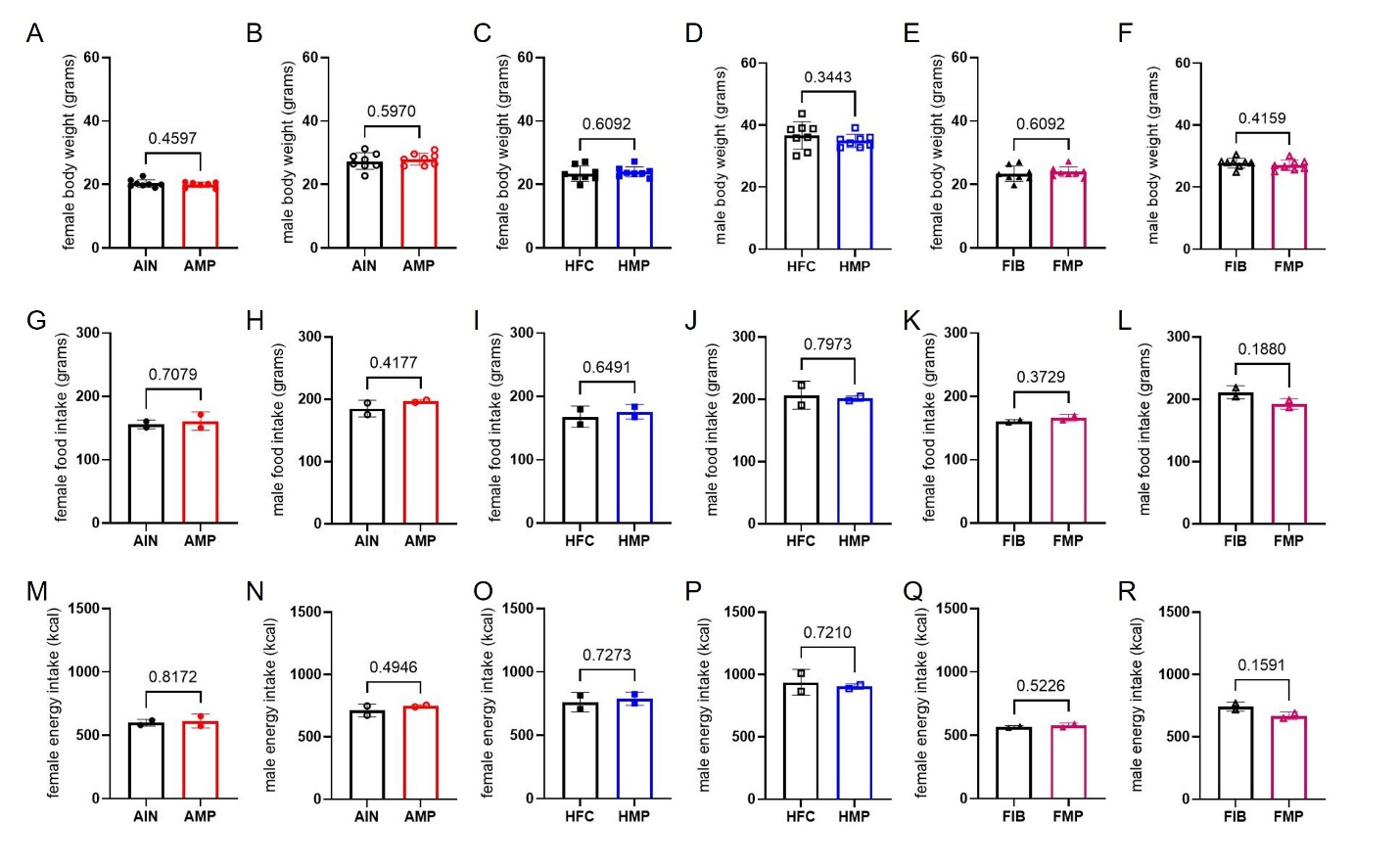
**Supplemental Figure 1. Microplastic intake regardless of diet and did not affect body mass in female or male C57BL/6J mice**. (A-F), final body weights for basal (A, B), high-fat (C, D), and high-fiber (E, F) diets. (G-L), total food intake for basal (G, H), high-fat (I, J), and high-fiber (K, L) diets. (M-R), total energy intake for basal (M, N), high-fat (O, P), and high-fiber (Q, R) diets. Open circles, squares and triangles represent female mice whereas closed circles, squares and triangles represent male mice. Data are mean ± SD; dots represent individual mice (n=8) or cage and analyzed using two-tailed unpaired Student’s t test or Welch’s t test.

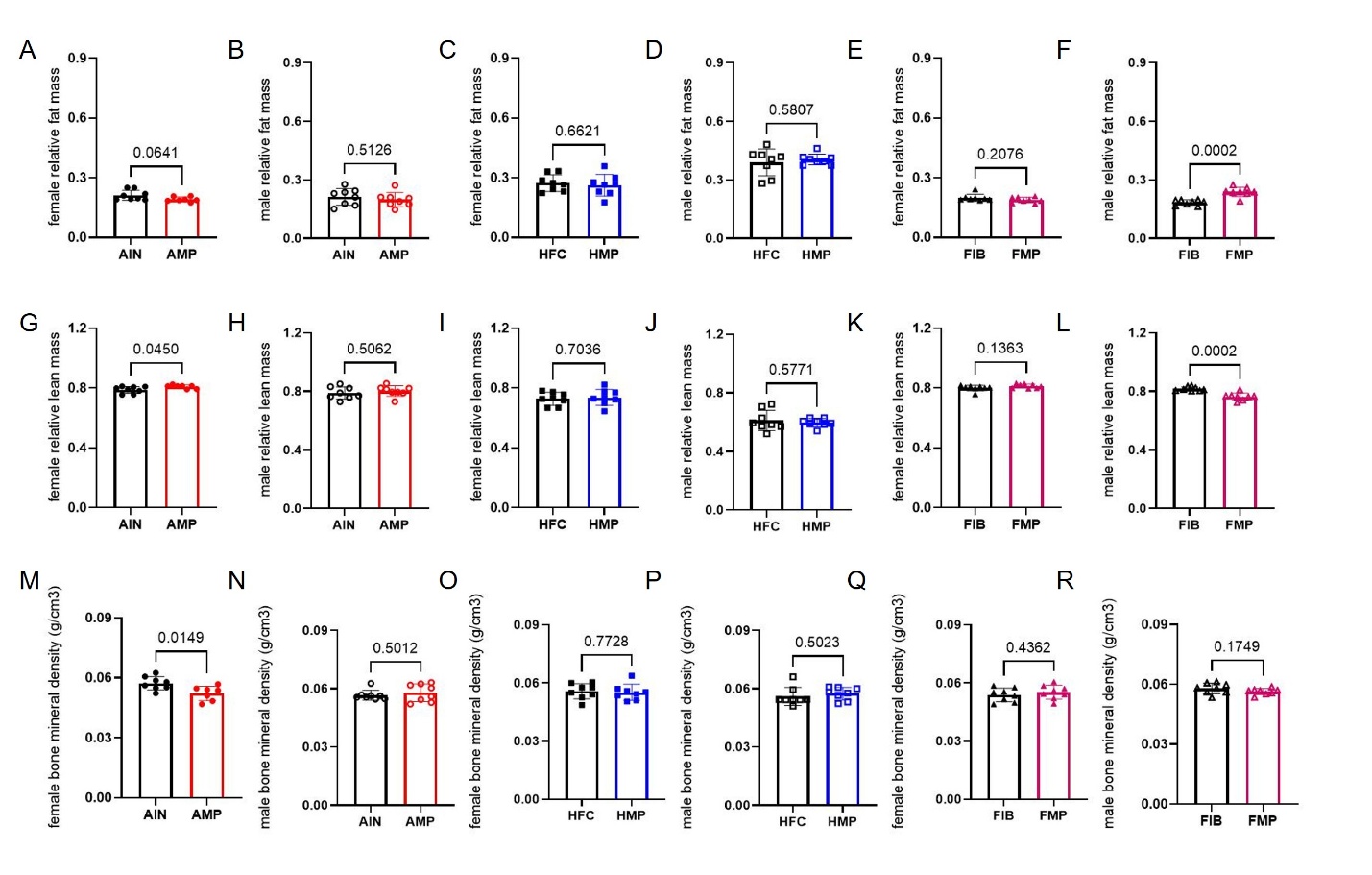
**Supplemental Figure 2. Sex- and diet-dependent alterations following microplastic exposure in body composition and bone mineral density assessed by DEXA at 12 weeks**. (A-F), terminal fat mass for basal (A, B), high-fat (C, D), and high-fiber (E, F) diets. (G-L), terminal lean mass for basal (G, H), high-fat (I, J), and high-fiber (K, L) diets. (M-R), terminal bone mineral density for basal (M, N), high-fat (O, P), and high-fiber (Q, R) diets. Open circles, squares and triangles represent female mice whereas closed circles, squares and triangles represent male mice. Data are mean ± SD; dots represent individual mice and analyzed using two-tailed unpaired Student’s t test or a Welch’s t test.

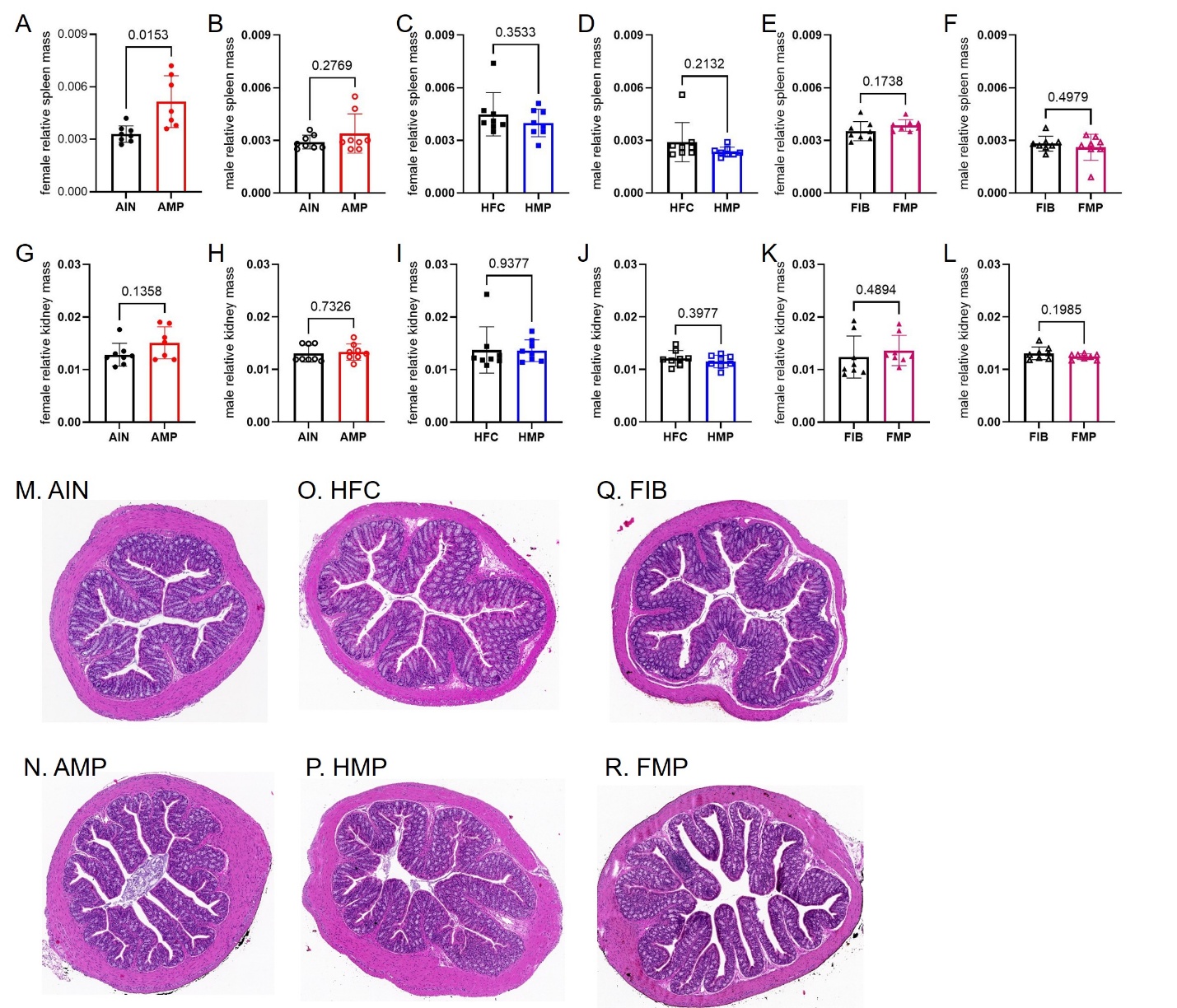
**Supplemental Figure 3. Dietary microplastic exposure, organ size and colonic histology. (A-F), Spleen mass in male and female mice in AIN and AMP groups (A, B); in HFC and HMP groups (C, D); and in FIB and FMP groups (E, F). (G-L), (A-F), Kidnet mass in male and female mice in AIN and AMP groups (G, H); in HFC and HMP groups (I, J); and in FIB and FMP groups (K, L).** Closed circles, squares and triangles represent female mice whereas opened circles, squares and triangles represent male mice. **(M-R),** Representative H&E-stained colon sections from mice on a control diet, AIN (M), control diet with MPs, AMP (N), high fat/high cholesterol diet, HFC (O), high fat/high cholesterol diet with MPs, HMP (P), high fiber diet, FIB (Q), and high fiber diet with MPs, FMP (R). Data are mean ± SD; dots represent individual mice and analyzed using two-tailed unpaired Student’s t test or a Welch’s t test.

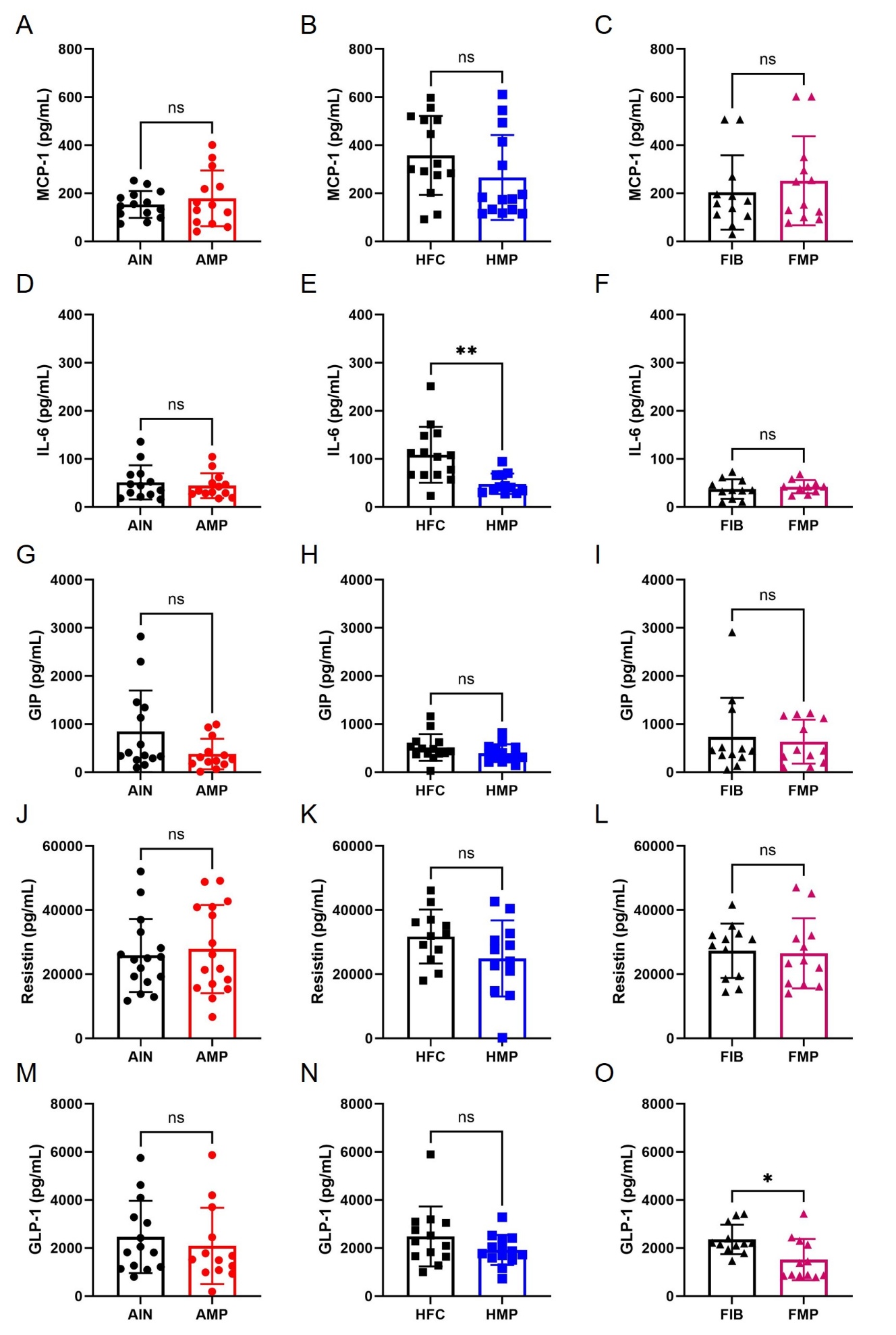
**Supplemental Figure 4. Dietary microplastic exposure does not cause broad hormonal changes in the plasma of mice. (A-O),** Circulating cytokines and hormones were assessed in plasma from mice on a control diet (AIN), control diet with MPs (AMP), high fat/high cholesterol diet (HFC), high fat/high cholesterol diet with MPs (HMP), high fiber diet (FIB), and high fiber diet with MPs (FMP). (A-C) MCP-1; (D-F) IL-6; (G-I) GIP; (J-L) Resistin; (M-O) GLP-1. Data are mean ± SD; dots represent individual mice. Data analyzed using Student’s t-test. Statistical significance is indicated as *P < 0.05; ns, not significant.

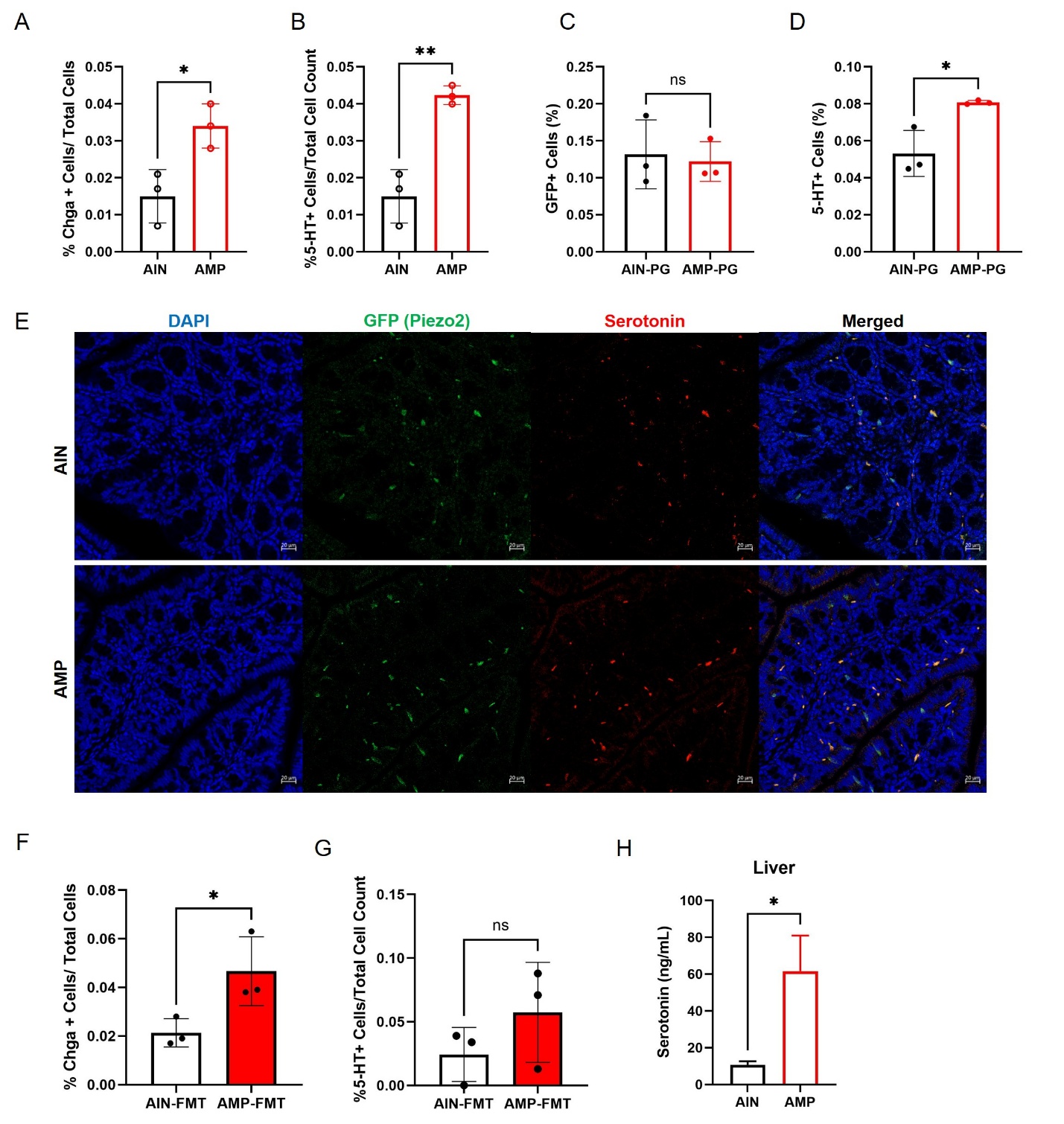
**Supplemental Figure 5. Colonic enteroendocrine cell abundance in Piezo2-GFP mice and after fecal microbiota transplant.** (A-B), Quantification of colonic enteroendocrine cells by immunofluorescence demonstrating increased proportions of CHGA^+^ cells (A) and 5-HT^+^ cells (B) in female AMP-fed mice relative to AIN female controls. (C-D), Quantification of GFP^+^ enterochromaffin cell abundance (C) and serotonin immunoreactivity per GFP^+^ cell (D) in colonic epithelium from AIN- and AMP-fed Piezo2-GFP mice. (E) Representative immunofluorescence images of proximal colon from AIN- and AMP-fed Piezo2-GFP reporter mice, stained for nuclei (DAPI, blue), GFP (green), and serotonin (5-HT, red). Merged images are shown. Scale bars, 20 μm. (F) Quantification of colonic enteroendocrine cells by immunofluorescence demonstrating increased proportions of CHGA^+^ cells (G) and no change in 5-HT^+^ cells (H) in recipient mice receiving fecal microbiota transplant from either AIN collected samples (AIN-FMT) or AMP collected samples (AMP-FMT). (H) Serotonin levels from the liver tissue homogenate from AMP-fed mice compared with AIN controls. Data are shown as mean ± s.d.; dots represent individual mice and analyzed using two-tailed unpaired Student’s t test. Statistical significance is indicated as *P < 0.05, **P < 0.01; ns, not significant.

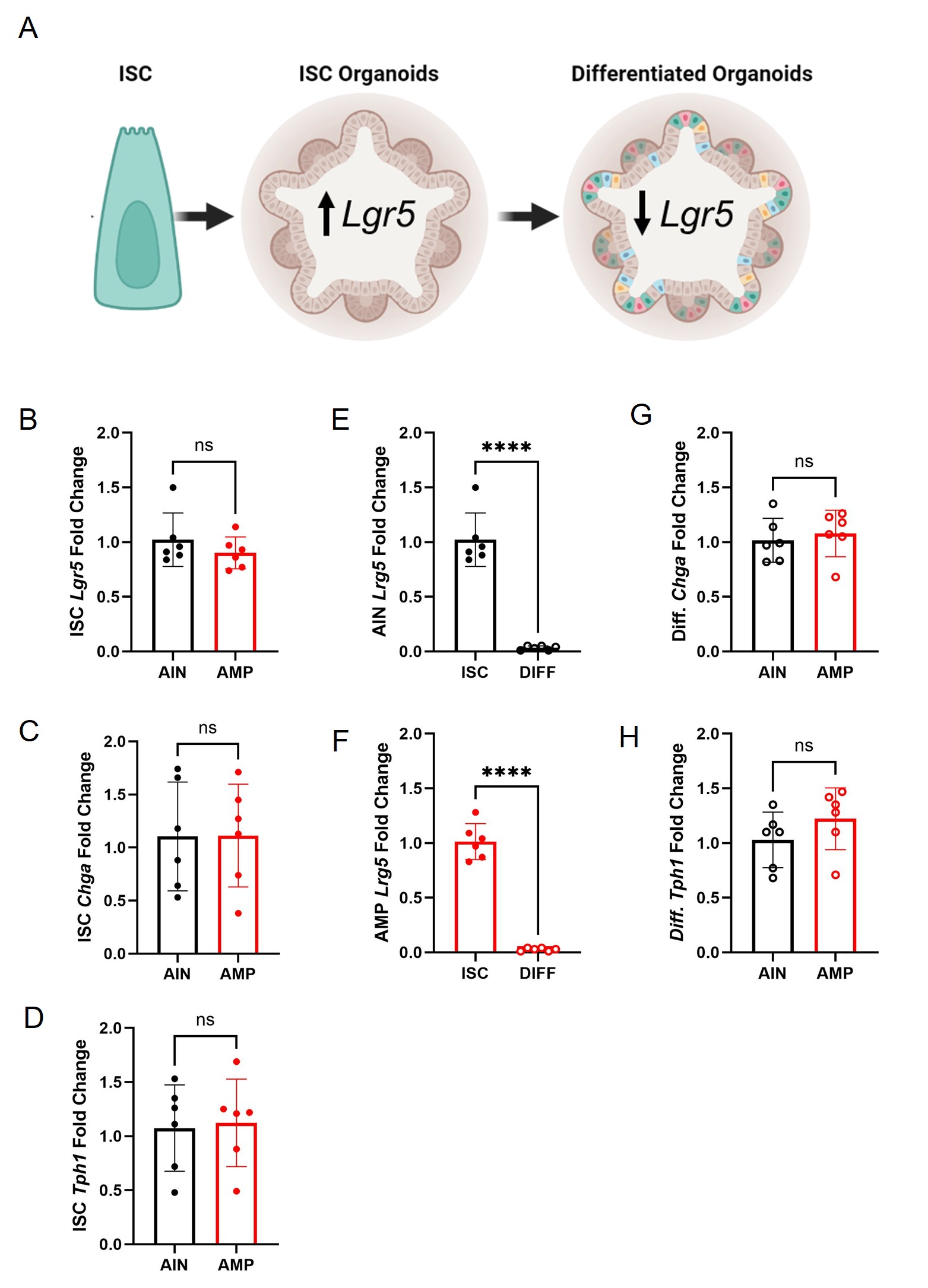
**Supplemental Figure 6. Colonic organoid lineage examination. (A), Cartoon showing intestinal stem cell to generate intestinal stem cell (ISC) organoids with high levels of *Lgr5* expression that can generate differentiated organoids with low levels of *Lgr5* expression. (B), *Lgr5*, an intestinal stem cell marker, expression in colonic organoids in a stem state comparing AIN- and AMP-derived colonic organoids. (C), *Chga* (chromogranin A), a pan enteroendocrine cell marker assessed** at a stem state in AIN- and AMP-**derived colonic organoids. (D), *Tph1* (**Tryptophan Hydroxylase 1), expressed in enterochromaffin cells, **assessed** at a stem state in AIN- and AMP-**derived colonic organoids. (E-F), Comparison of *Lgr5* expression in colonic organoids in a stem state versus differentiated state in AIN- (E) or AMP-derived colonic organoids (F). (G), *Chga* expression in** AIN- and AMP-**derived differentiated colonic organoids. (H), *Tph1* expression in** AIN- and AMP-**derived differentiated colonic organoids.** Data are shown as mean ± s.d.; dots represent individual mice. Statistical significance is indicated as ****P < 0.001; ns, not significant.

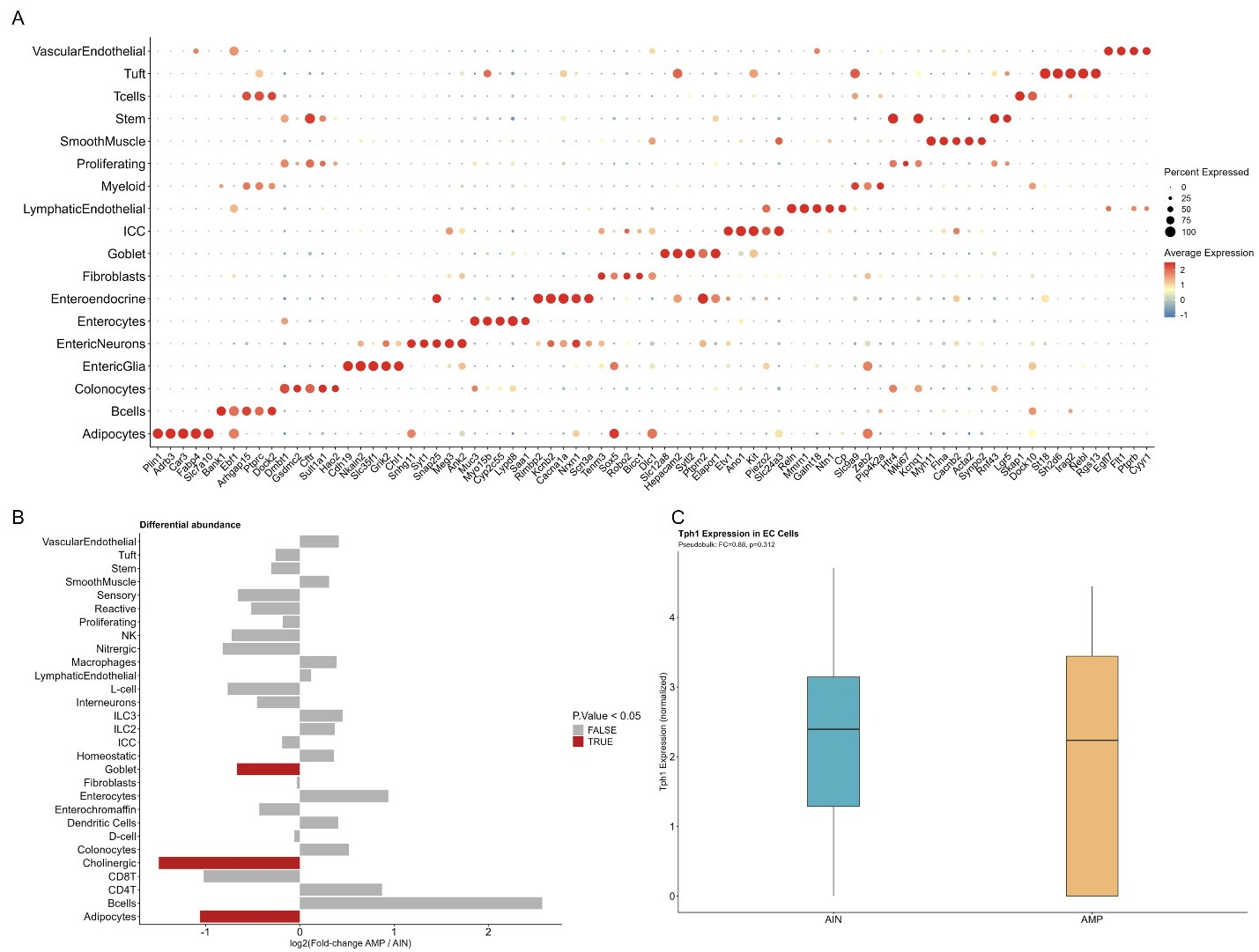
**Supplemental Figure 7. Single nuclei RNA sequencing and cell specific transcriptional expression. (A), Dot plot depicting mean expression and percent-expressing cells across all different cells types found within the colon indicating the highest expressed markers per cell type. (B), Differential abundance through log2(fold change) where goblet, cholinergic, and adipocytes were more concentrated in microplastic treated colons (AMP) as compared to the control (AIN). (C), Normalized *Tph1* expression on enterochromaffin cells in AIN and AMP colons.** n=4 per group.

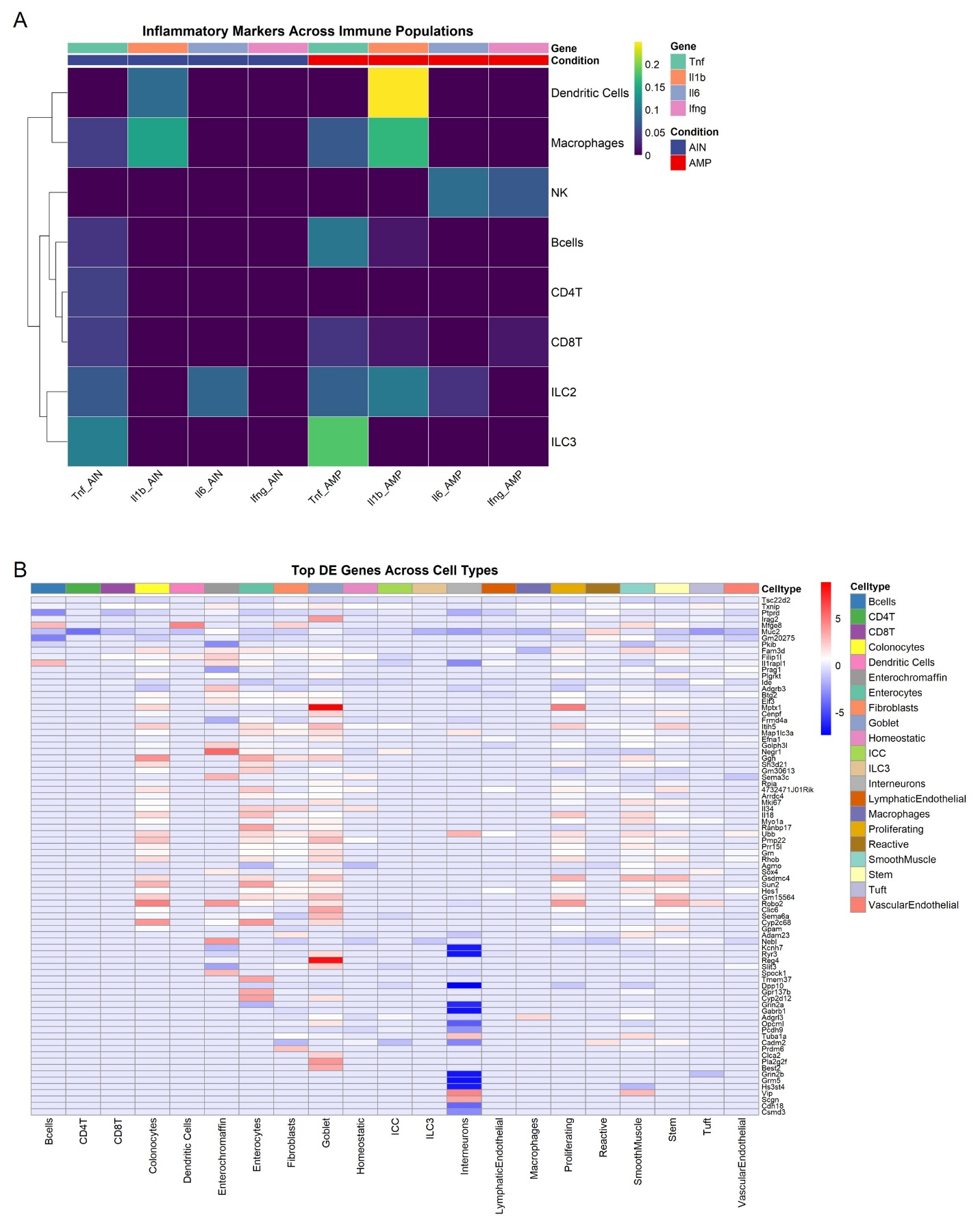

**Supplemental Figure 8. Single nuceli RNA sequencing and inflammatory cell specific transcriptional expression. (A), heat map evaluating inflammatory markers across immune populations. (B), heat map displaying the top differentially expressed genes across all cell types.** n=4 per group.

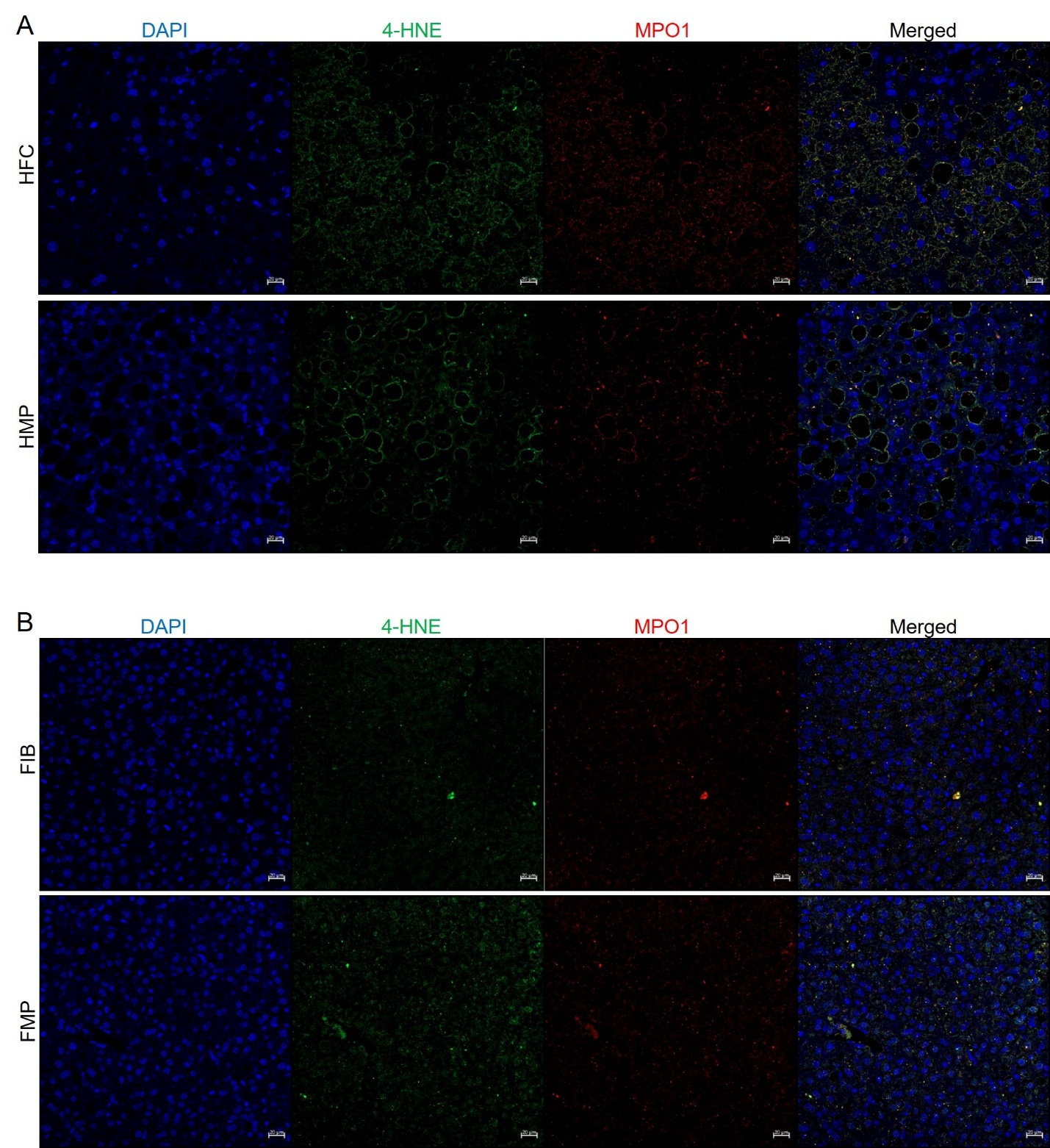
**Supplemental Figure 9. Representative immunofluorescence detection of hepatic oxidative stress in other dietary conditions mice.** (A-B) Representative liver sections from HFC and HMP (A) or FIB and FMP (B) mice stained for 4-hydroxynonenal (4-HNE; green) and myeloperoxidase (MPO1; red), and nuclei counterstained with DAPI (blue). Merged images are shown in the far-right panels. Scale bars, 20 μm.

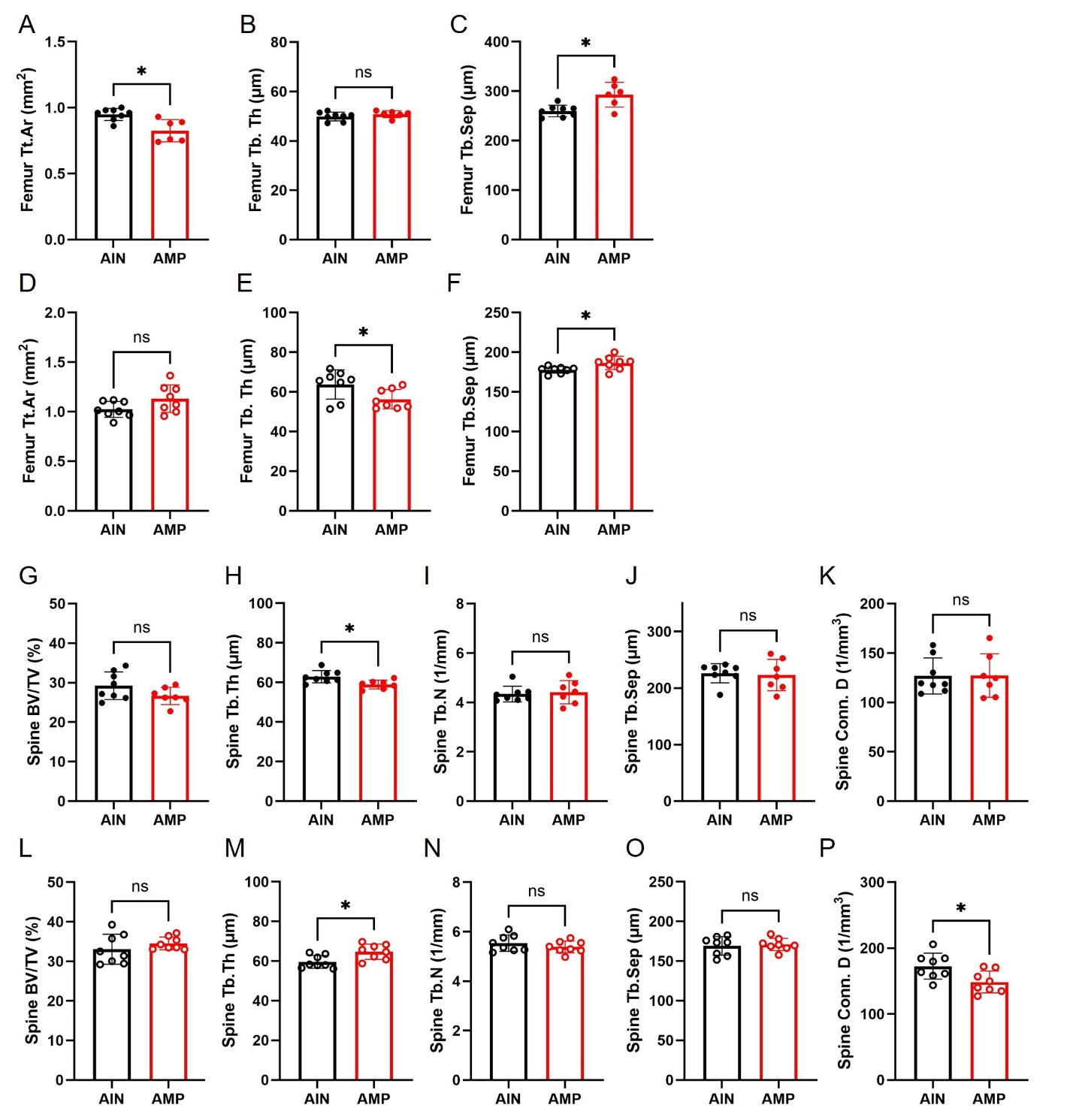
**Supplemental Figure 10. Dietary microplastic exposure disrupts vertebral bone microarchitecture in a sex-specific manner.** (A-C), Femoral µCT analysis in females (closed circles) showing reduced (A) total area (Tt.Ar), (B) trabecular thickness (Tb.Th), (**C**) with increased trabecular separation (Tb.Sep). (D-F), Femoral µCT analysis in males (open circles) showing (D) total area (Tt.Ar), (E) decreased trabecular thickness (Tb.Th), (**F**) with increased trabecular separation (Tb.Sep) in AMP-fed mice relative to AIN controls. (G-K), Spinal µCT analysis in females (closed circles) demonstrating preserved vertebral parameters (**G, I-K**) but reduced spinal trabecular thickness (Tb.Th) (**H**), in AMP-fed mice. **(L-P),** Spinal µCT analysis in males (open circles) demonstrating preserved vertebral parameters (**L, N, O**) but increased spinal trabecular thickness (Tb.Th) (M) and reduced spinal connective density (Conn.D) (P), in AMP-fed mice. Closed circles represent female mice whereas opened circles represent male mice. Data are mean ± SD; dots represent individual mice. Statistical significance is indicated as *P < 0.05; ns, not significant.

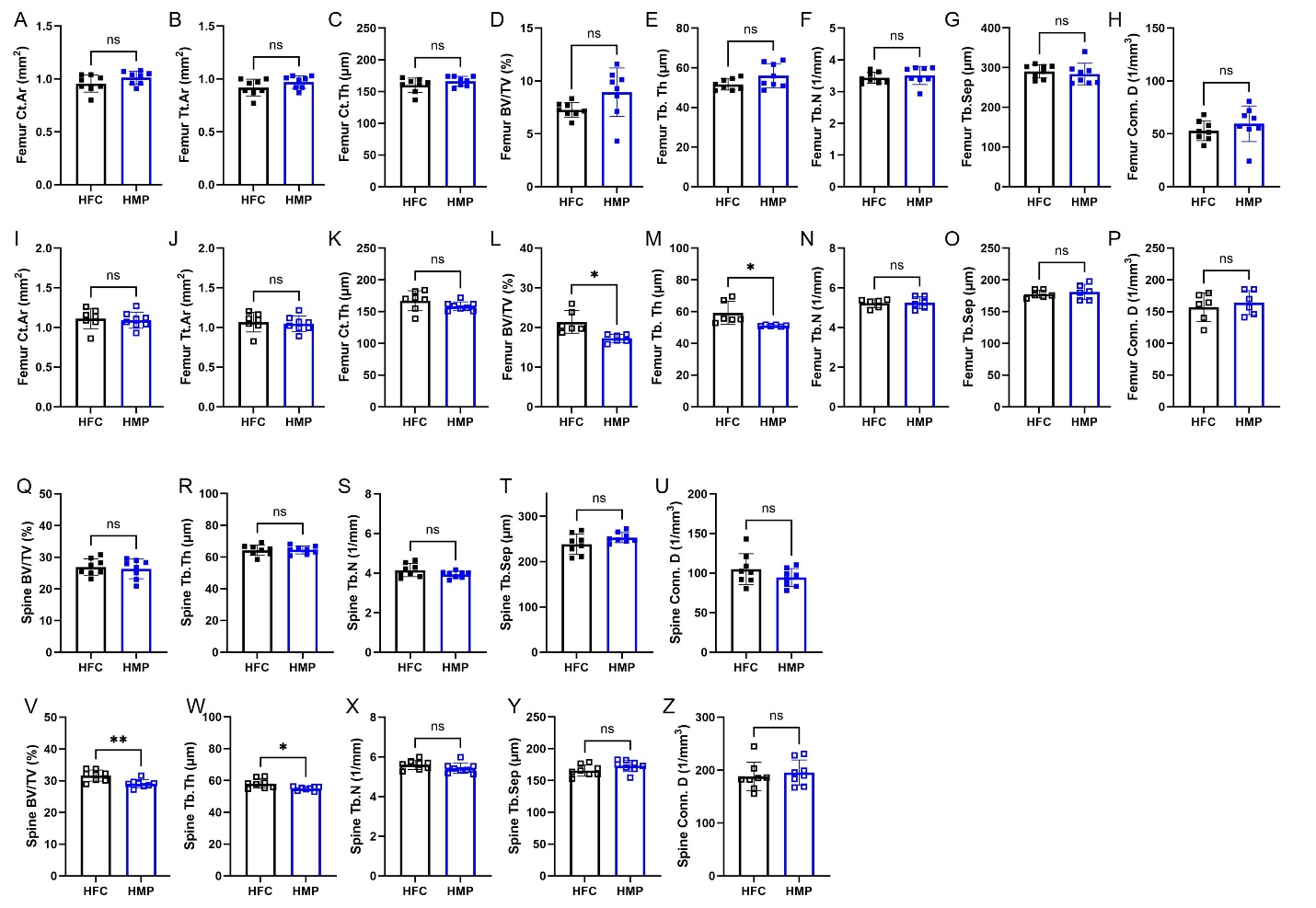
**Supplemental Figure 11. Dietary microplastic exposure disrupts bone microarchitecture in male mice on a high fat diet.** (A-H), Femoral µCT analysis in females (closed squares) showing preserved vertebral parameters (Ct.Ar; Tt.Ar; Ct.Th;BV/TV; Tb.Th; Tb.N; Tb.Sep; and Conn.D) in HMP-fed mice relative to HFC controls. (I-P), Femoral µCT analysis in males (open squares) demonstrating preserved vertebral parameters (Ct.Ar; Tt.Ar; Ct.Th;BV/TV; Tb.Th; Tb.N; Tb.Sep; and Conn.D), (**I-P**) but reduced femur bone volume (BV/TV) (**L)**, and trabecular thickness (Tb.Th) (**M**) in HMP-fed mice. (Q-U), Spinal µCT analysis in females (closed squares) demonstrating preserved vertebral parameters (BV/TV; Tb.Th; Tb.N; Tb.Sep; and Conn.D) in HMP-fed mice compared to HFC-fed mice. (V-Z), Spinal µCT analysis in males (open squares) demonstrating preserved vertebral parameters (BV/TV; Tb.Th; Tb.N; Tb.Sep; and Conn.D), (**X-Z**) but decreased spinal bone volume (BV/TV) (v) and trabecular thickness (Tb.Th) (W), in HMP-fed mice. Closed squares represent female mice whereas opened squares represent male mice. Data are mean ± SD; dots represent individual mice. Statistical significance is indicated as *P < 0.05, **P < 0.01; ns, not significant.

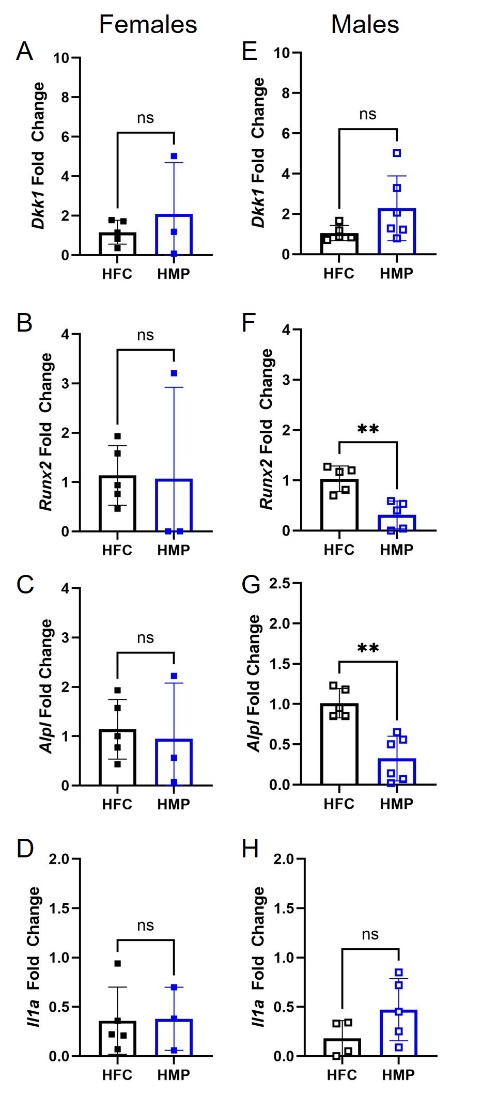

**Supplemental Figure 12. Dietary microplastic exposure alters transcriptional response in the bone of male mice on a high fat diet.**  (A-D), Relative mRNA expression of Dkk1 (**A**), Runx2 (**B**), Alpl (**C**), and Il1a (**D**) in femoral bone from females. (E-H), Relative mRNA expression of Dkk1 (**E**), Runx2 (**F**), Alpl (**G**), and Il1a (**H**) in femoral bone from males. Closed squares represent female mice whereas opened squares represent male mice. Data are mean ± SD.; dots represent individual mice. Statistical significance is indicated as **P < 0.01; ns, not significant.

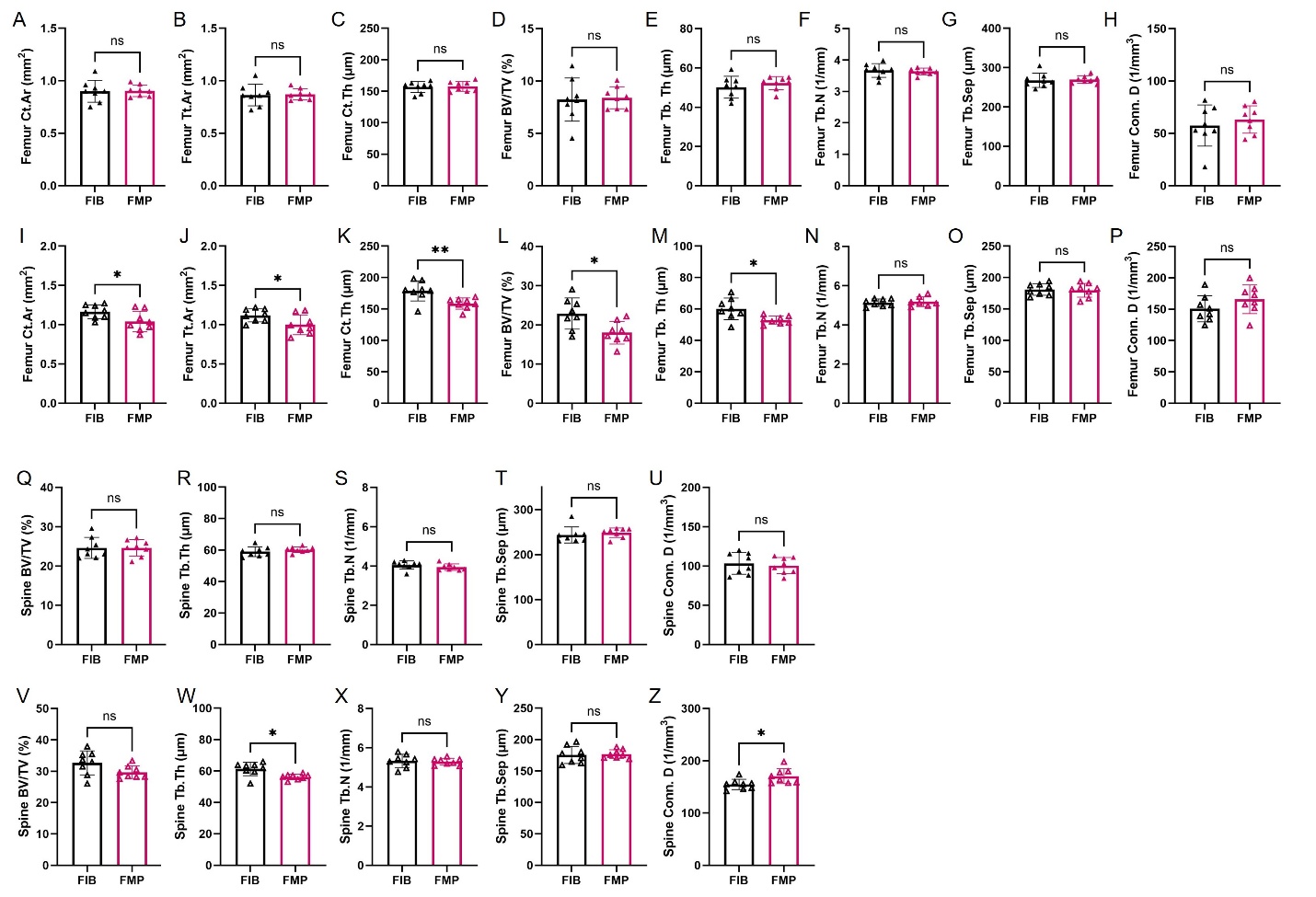
**Supplemental Figure 13. Dietary microplastic exposure disrupts bone microarchitecture in male mice on a high fiber diet.** (A-H), Femoral µCT analysis in females (closed triangles) showing preserved vertebral parameters (Ct.Ar; Tt.Ar; Ct.Th;BV/TV; Tb.Th; Tb.N; Tb.Sep; and Conn.D) in FMP-fed mice relative to FIB controls. (I-P), Femoral µCT analysis in males (open triangles) demonstrating preserved vertebral parameters (Ct.Ar; Tt.Ar; Ct.Th;BV/TV; Tb.Th; Tb.N; Tb.Sep; and Conn.D), (**N-P**), but reduced femur cortical area (Ct.Ar) (**I)**, total area (Tt.Ar) (**J**), cortical thickness (Ct.Th) (K), bone volume (BV/TV) (L), and trabecular thickness (Tb.Th) (M) in FMP-fed mice. (Q-U), Spinal µCT analysis in females (closed triangles) demonstrating preserved vertebral parameters (BV/TV; Tb.Th; Tb.N; Tb.Sep; and Conn.D) in FMP-fed mice compared to FIB-fed mice. (V-Z), Spinal µCT analysis in males (open triangles) demonstrating preserved vertebral parameters (BV/TV; Tb.Th; Tb.N; Tb.Sep; and Conn.D), (**V, X-Y**) but decreased spinal trabecular thickness (Tb.Th) (W) and bone connective density (Conn.D) (Z), in FMP-fed mice. Closed triangles represent female mice whereas opened triangles represent male mice. Data are mean ± SD; dots represent individual mice. Statistical significance is indicated as *P < 0.05, **P < 0.01; ns, not significant.

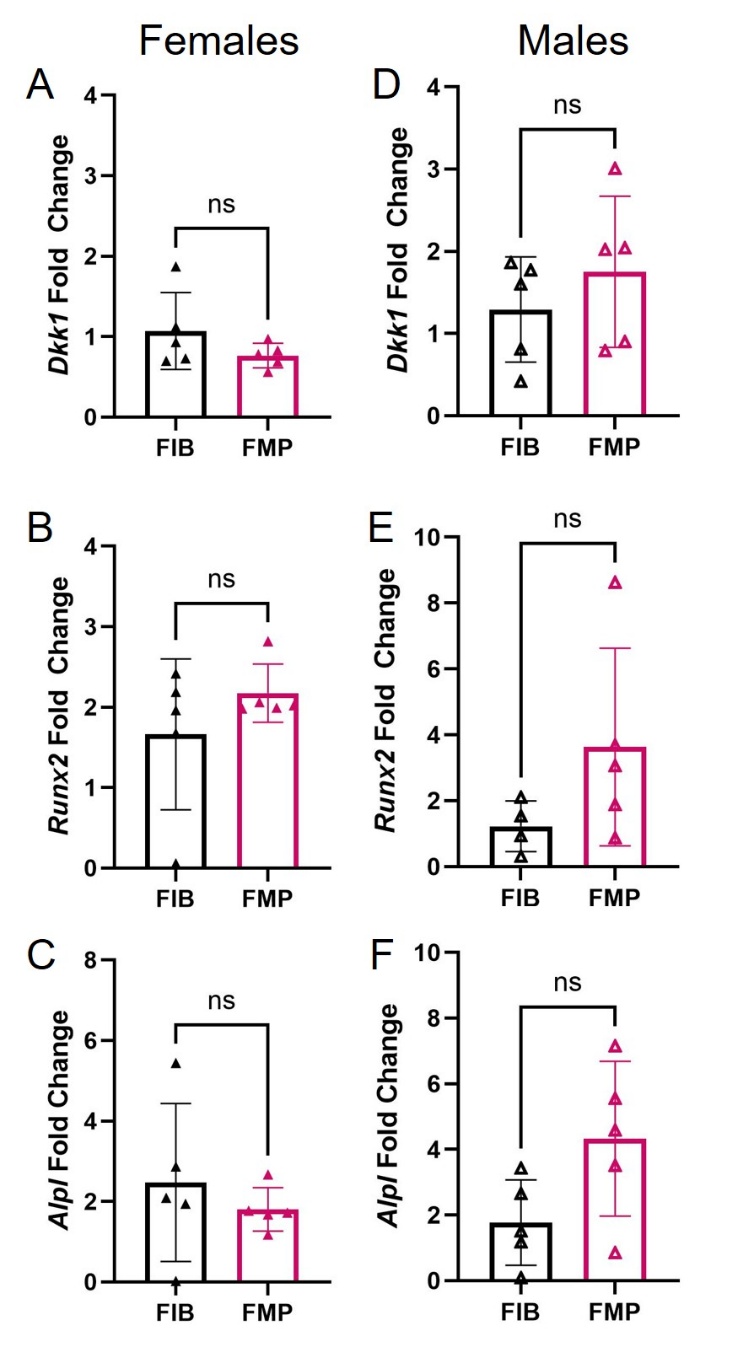

**Supplemental Figure 14. Dietary microplastic exposure do not alter transcriptional response in the bone of mice on a high fiber diet.**  (A-C), Relative mRNA expression of Dkk1 (**A**), Runx2 (**B**), Alpl (**C**) in femoral bone from females. (D-F), Relative mRNA expression of Dkk1 (**D**), Runx2 (**E**), Alpl (**F**), and in femoral bone from males. Closed triangles represent female mice whereas opened triangles represent male mice. Data are mean ± SD; dots represent individual mice. Statistical significance is indicated as ns, not significant.

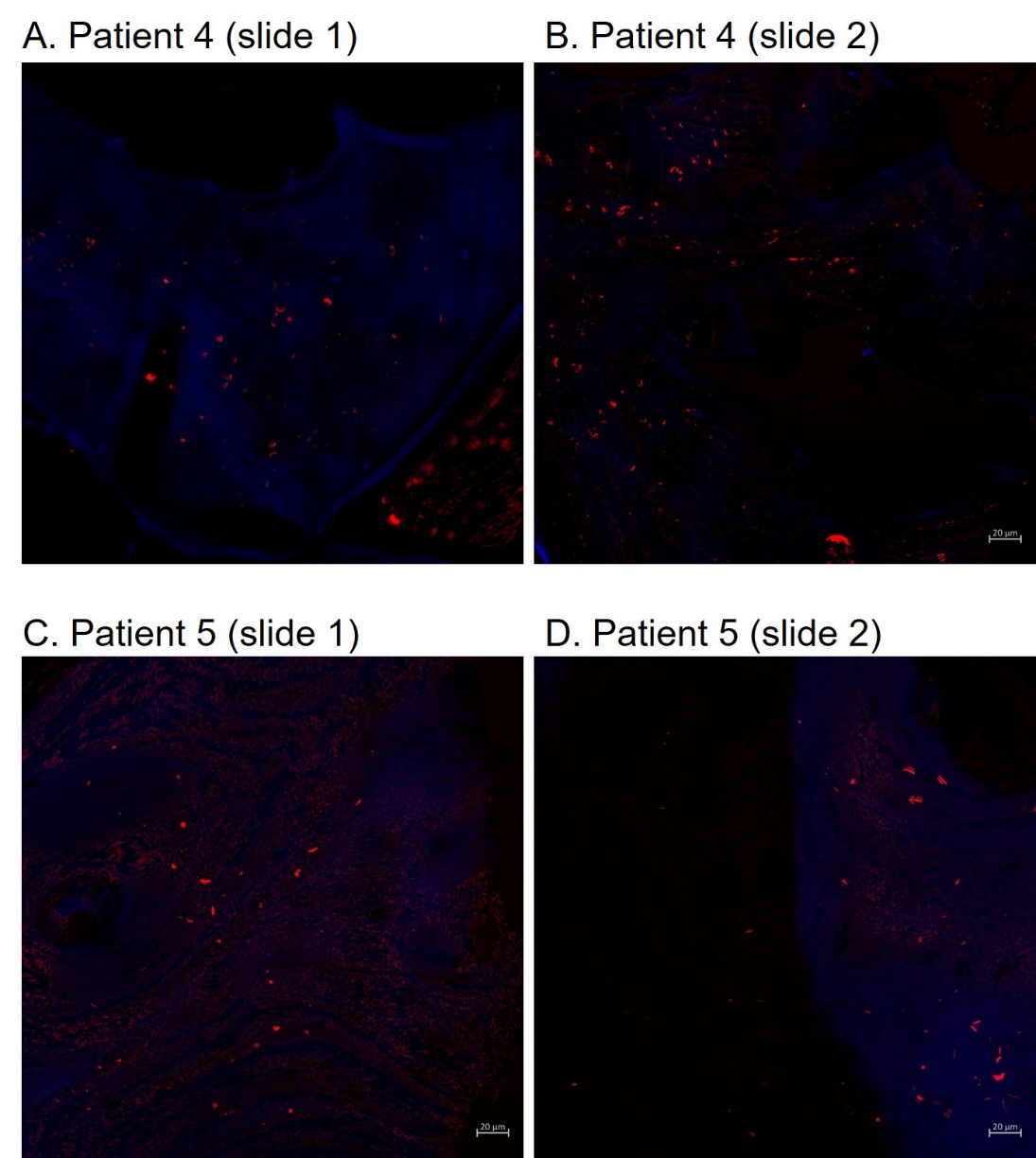

**Supplemental Figure 15. Polymer detection in human distal femur tissue from donors without orthopedic prostheses.** (A-D) Fluorescence images of histological sections from distal femurs from two of the independent human donors (Patients 4-5; donor information in Supplemental Table 6) stained a conjugated polymer nanoparticle-based MP dye (MP-DYE, red) and control nanoparticle-based dye (blue). Demonstrating discrete polymer-positive signals within intact bone tissue. Objective magnification: Scale bars, 20 µm.

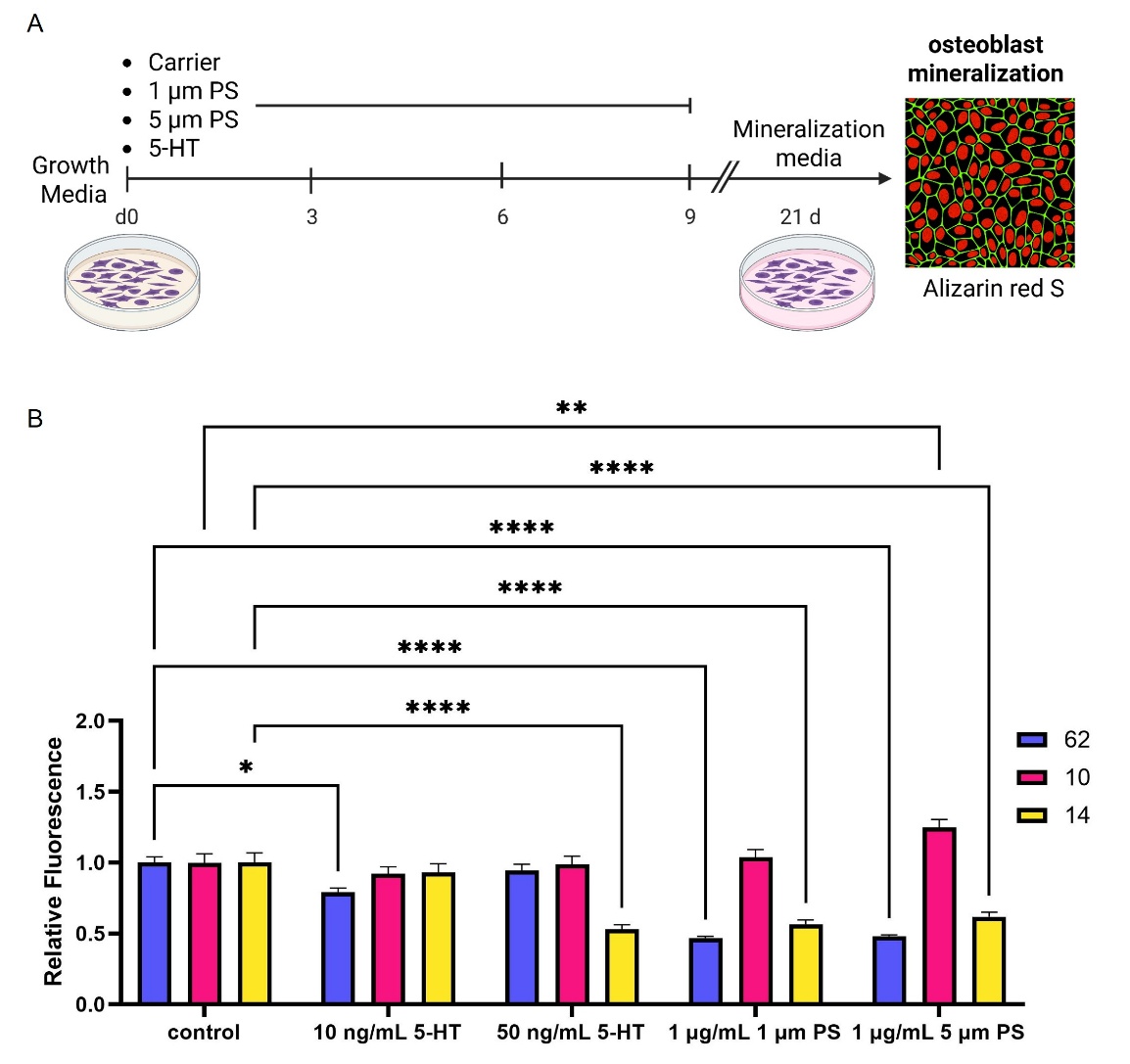

**Supplemental Figure 16. Inter-individual variability in osteoblast mineralization following polystyrene microsphere and serotonin exposure**. (A), ***in vitro* approach where t**hree human osteoblast cell lines (identifiers 62, 10, and 14 are individual donors, donor information can be found in Supplemental Table 8) were treated with either 10 ng/mL of 5-HT, 50 ng/mL of 5-HT, 1 µg/mL of 1 µm PS, 1 µg/mL of 5 µm PS, or vehicle control for 10 days (day 0-9) in Osteoblast Growth Medium followed by 21 days is Osteoblast Mineralization Medium, then stained with Alizarin Red S and DAPI. (B), Alizarin:DAPI fluorescence ratios for each treatment group were normalized to DMSO controls; data are shown as mean ± SEM and analyzed using a two-way ANOVA with Tukey’s multiple comparison test. Statistical significance is indicated as *P < 0.05, **P < 0.01, ****P < 0.001.
